# Autophagy modulates apical growth and development in the moss *Physcomitrium patens*

**DOI:** 10.1101/2022.04.19.488653

**Authors:** Georgina Pettinari, Juan Finello, Macarena Plaza Rojas, Franco Liberatore, Germán Robert, Santiago Otaiza-González, Pilar Velez, Martin Theumer, Patricia Agudelo-Romero, Claudio Gonzalez, Ramiro Lascano, Laura Saavedra

**Affiliations:** Unidad Ejecutora de Doble Dependencia INTA-CONICET (UDEA), Córdoba, Argentina; Cátedra de Fisiología Vegetal, Facultad de Ciencias Exactas, Físicas y Naturales, Universidad Nacional de Córdoba; Córdoba, Argentina; Instituto de Investigación Médica Mercedes y Martín Ferreyra - INIMEC (CONICET) - Universidad Nacional de Córdoba, Córdoba, Argentina.; Centro de Investigaciones en Bioquímica Clínica e Inmunología (CIBICI), Universidad Nacional de Córdoba-CONICET, Córdoba, Argentina; Telethon Kids Institute, Perth Children’s Hospital, Nedlands WA 6009, Australia

**Keywords:** Apical growth, ATG8, Autophagy, Bryophytes, Carbon starvation, Cell death, Chloronema and caulonema, Development, Nitrogen starvation, Senescence

## Abstract

Different to root hairs and pollen tubes, *Physcomitrium patens* apical growing protonemal cells have the singularity that they continue to undergo cell divisions as the plant develops, allowing to study autophagy in the context of a multicellular apical growing tissue coupled to development. Herein, we showed that the core autophagy machinery is present in the moss *P. patens*, and deeply characterized the growth and development of wild-type, *atg5* and *atg7 loss-of-function* mutants under optimal and nutrient-deprived conditions. Our results showed that the growth of the different morphological and functional protonemata apical growing cells, chloronema and caulonema, is differentially modulated by this process. These differences depend on the protonema cell type and position along the protonemal filament, and growth condition. As a global plant response, the absence of autophagy triggers the spread of the colony through protonemata growth at the expense of a reduction in buds and gametophore development, and thus the adult gametophytic and reproductive phases. Altogether this study provides valuable information indicating that autophagy has roles during apical growth with differential responses within the cell types of the same tissue and contributes to life cycle progression and thus the development of the 2D and 3D tissues of *P. patens*.

**HIGHLIGHT:** Autophagy is differentially induced in protonemal cells, and contributes to apical growth, life cycle progression, and thus the development of the 2D and 3D tissues of *P. patens*.

## INTRODUCTION

Macroautophagy (hereafter autophagy) is an essential catabolic pathway for eukaryotic cell homeodynamics that mediates cellular recycling during development and stress conditions (Marshall and Vierstra, 2018), involving the coordinated interaction among more than 30 highly conserved autophagy-related (*ATG*) proteins and components of the secretory system (Farhan *et al*., 2017; Nakatogawa, 2020; Soto-Burgos *et al*., 2018; Wang *et al*., 2020). During this process, a wide range of intracellular material is engulfed into double-membrane structures termed autophagosomes, and transported this autophagic cargo to vacuoles in plant and yeast cells (or lysosomes in mammalian cells), to be degraded and recycled for different cellular purposes or remobilized to other parts of the organism (Marshall and Vierstra, 2018).

Most knowledge in the field of plant autophagy arises from studies in Angiosperms, mainly in the model *Arabidopsis thaliana,* or crops such as *Zea mays*, *Oryza sativa,* and *Glycine max* (Tang and Bassham, 2018), pointing out that autophagy is virtually involved in all aspects of plant physiology (Avin-Wittenberg *et al*., 2015; Di Berardino *et al*., 2018; Doelling *et al*., 2002; Estrada-Navarrete *et al*., 2016; Guiboileau *et al*., 2012; Hanaoka *et al*., 2002; Kwon *et al*., 2010; Li *et al*., 2015; Li *et al*., 2020; Sera *et al*., 2019). Land plants (embryophytes) are a monophyletic lineage that evolved from within freshwater streptophyte algae c. 470–515 mya (Puttick *et al*., 2018). The recent availability of sequenced genomes from several lineages of streptophyte algae, and bryophyte species, the first group of plants that conquered land, has allowed comprehensive comparative evolutionary analyses moving forward our understanding of plant evolution (Rensing, 2017, 2020). Studies on autophagy in the algae *Chlamydomonas reinhardtii* (Kajikawa and Fukuzawa, 2020; Pérez-Pérez *et al*., 2012; Pérez-Pérez *et al*., 2017), the liverwort *Marchantia polymorpha* (Norizuki *et al*., 2019), and the moss *Physcomitrium (Physcomitrella) patens* (Chen *et al*., 2020; Kanne *et al*., 2021; Mukae *et al*., 2015; Sanchez-Vera *et al*., 2017) have started to emerge, providing progress on evolutionary aspects of plant autophagy as well as species-specific roles of this process.

*P. patens* has been well established as a model system (Rensing *et al*., 2020). It is also a model for polar growth studies because similar to unicellular root hairs and pollen tubes, rhizoids and protonemata exhibit apical growth (Rounds and Bezanilla, 2013; Vidali and Bezanilla, 2012), but in contrast to them, have the singularity that continues to undergo cell divisions as the plant develops, allowing to study this process in the context of a multicellular tissue (Rounds and Bezanilla, 2013). Protonemal cells live longer than pollen tubes and root hairs and integrate considerably more environmental signals during their growth and development (Bascom *et al*., 2018). Protonemata is composed of two morphologically and functionally distinct cells: chloronemata and caulonemata. Whereas chloronemata are rich in chloroplasts, divide every 24 hours, and have a photosynthetic role, caulonemata contain fewer and less developed chloroplasts, divide every 8 hours, and have a role in substrate colonization and nutrient acquisition (Menand *et al*., 2007a). Caulonema differentiates from a chloronemal tip cell, and this transition is regulated by auxin levels and energy availability (Thelander *et al*., 2018; Thelander *et al*., 2005; Viaene *et al*., 2014). Thus, caulonemata growth is favored by high light conditions or an external carbon source such as glucose (Thelander *et al*., 2005), but also by nutrient deficiencies such as phosphate starvation (Wang *et al*., 2008) or growth with nitrate as a single nitrogen source, probably due to a carbon/nitrogen unbalance.

Based on these facts, our questions were raised on whether autophagy plays a role in regulating moss protonemata nutritional status and thus, apical growth and development in the context of this multicellular tissue. It is unknown whether protonemata cells have differential responsiveness to canonical autophagy-inducing conditions such as carbon and nitrogen deficiencies, or whether autophag`y plays a role not only under these deficiencies but also under the dark period of a long-day (LD) in optimal growth conditions.

In this study, we described the ATG system in *P. patens* and explored the autophagic response of wild-type, *atg5*, and *atg7* loss-of-function lines by phenotypic characterization, and gene expression analysis, combined with autophagic flux assays and visualization of autophagic vesicles using a *PpATG8b::GFP-PpATGb* reporter line. Our results showed that autophagy highly contributes to *P. patens* apical growth and development, under both optimal and nutrient-deprived conditions, and suggested that autophagy mainly acts as a mechanism that provides energy to sustain apical growth coupled to development.

## MATERIALS AND METHODS

### Identification of *ATG* genes in *P. patens* and phylogenetic analysis

Orthologs *ATG* genes from the moss *P. patens* were retrieved by reciprocal Basic Local Alignment Search Tool for Protein (BLASTP) and Nucleotide (TBLASTN) searches using the coding regions of *ATG* genes previously described in *A. thaliana* performed in Phytozome v12.1 database (https://phyto zome.jgi.doe.gov) (Table S1). For phylogenetic analyses, a similar search was performed for ATG8 protein sequences from several algae and plant species in Phytozome v12.1 database and ONEKP: BLAST for 1,000 Plants (https://db.cngb.org/onekp/species/) (Table S2). Evolutionary analyses were conducted in MEGAX (Kumar *et al*., 2018). Multiple sequence alignment was inferred using MUSCLE. The evolutionary history was inferred by using the Maximum Likelihood method and JTT matrix-based model and the bootstrap consensus tree was inferred from 1000 replicates.

### *PpATG8s* promoter and network analysis

Analysis of transcription factor binding sites (TFBSs) was done using upstream sequence regions between 0.6 kb and 2.5 kb, depending on the flanking gene boundaries according to *P. patens* genome v3.3 (https://phytozome.jgi.doe.gov/pz/portal.html). These sequences were investigated with Homer v4.11.1 software (https://homer.ucsd.edu/homer/motif/), assessing 506 motifs from the Homer plant database against the six *PpATG8s* genes using the parameter “-find” from the homer2 “findMotifs.pl” script. Then, motif score > 9 were selected for downstream analyses (Table S4). Gene–motif interaction network was built representing the relationship between gene and known motif employing Cytoscape 3.8.2 software (Shannon *et al*., 2003). Co-expression network table was generated using “NetworkAnalyzer” tool from Cytoscape, keeping nodes with at least two degrees (Table S5).

### Plant material, growth conditions

All experiments described in this study, including the generation of *P. patens* mutant lines, were performed with *Physcomitrium (Physcomitrella) patens ssp. patens* (Hedwig) ecotype ‘Gransden 2004’. Plant cultures were grown axenically at 24°C under a long-day photoperiod (16-h light/8-h dark) with a photon flux of 60-80 μmol.m^2^.s^-1^. *P. patens* protonemal tissue was subcultured routinely at 7-day intervals on cellophane disks (AA packaging) overlaying Petri dishes (90 mm in diameter) that contained BCDAT medium [0.92 g/L di-ammonium tartrate (C_4_H_12_N_2_O_6_), 0.25 g/L MgSO_4_.7H_2_O, 1.01 g/L KNO_3_, 0.0125 g/L FeSO_4_.7H_2_O, 0.25 g/L KH2PO4 (pH 6.5), 0.147 g/L CaCl_2_.2H_2_O, and 0.001% Trace Element Solution (0.055 g/L CuSO_4_.5H_2_O, 0.055 g/L ZnSO_4_.7H_2_O, 0.614 g/L H_3_BO_3_, 0.389 g/L MnCl_2_.4H_2_O, 0.055 g/L CoCl_2_.6H_2_O, 0.028 g/L KI, 0.025 g/L Na_2_MoO_4_.2H_2_O)] and 7 g/L agar, or minimal media BCD where the di-ammonium tartrate from BCDAT media was omitted (Ashton and Cove, 1977).

### Generation of *atg5* and *atg7* knock out mutants in *P. patens*

Polyethylene glycol–mediated protoplast transformation was performed according to (Saavedra *et al*., 2015). In short, 6-day-old protonemata were treated with 0.5% Driselase (Sigma-Aldrich) in 8.5% w/v mannitol for 30 min, passed through a 100-mm sieve, incubated for 15 min at room temperature, and passed through a 50-mm sieve. The protoplasts of the final flow-through, which were washed twice in 8.5% mannitol, were ready for further use. Protoplasts were transformed at a concentration of 1.6·10^6^ protoplasts.ml^-1^. Each transformation consisted of 0.3 ml of protoplast suspension and 10-15 μg of linear DNA. To eliminate any episomal-resistant colonies, two rounds of selection were undertaken using the appropriate antibiotic. Transformations with knock-out vectors were performed with plasmid-based vectors linearized with the appropriate restriction enzyme (Thermo Fisher Scientific).

For colony experiments in response to starvation, moss colonies were initiated from 1 mm^2^ spot cultures on cellophane overlaid BCDAT media and grow for 14 days. Then the cellophane was transferred to new media for control (BCDAT), and then kept under optimal growth light conditions (LD photoperiod) either with nitrogen (LN) or without nitrogen supply (L-N), or transferred to darkness (DN) or darkness with an external supplement of sucrose (DN+S), for additional 3 or 7 days. For colony experiments in minimal media (BCD), moss colonies were initiated from 1 mm^2^ spot cultures on cellophane overlaid BCD media without or with additives applied to the media to a final concentration of 1 μM for auxin (1-Naphthylacetic acid, NAA, Sigma-Aldrich), 0.2 μM trans-Zeatin (tZ, Sigma-Aldrich) or 2% (w/v) Sucrose (Sigma-Aldrich). Moss colonies were imaged with an Olympus SZX16 stereomicroscope equipped with a color camera (Olympus DP71).

For protonemata carbon starvation experiments, protonemata were first grown for 7 days under LD photoperiod (16-h light/8-h dark) on solid BCDAT medium before being transferred to darkness without or supplemented with 2% sucrose during the indicated number of hours (h). For nitrogen starvation experiments, protonemata were first grown for 7 days under LD photoperiod (16-h light/8-h dark) on solid BCDAT medium containing 0.7% agar before transfer to media where KNO3 and ammonium tartrate from BCDAT medium was replaced with potassium sodium tartrate tetrahydrate. Experimental conditions for nitrogen starvation gene expression analysis and visualization of autophagic particles were performed under continuous light to discriminate the effects of carbon and nitrogen deficiencies.

For LY294002 (Sigma-Aldrich, stock solution diluted in DMSO) experiments, 40 μM of the drug was added to solid growth media before it solidified. The tissue was transferred to LY containing media at the time of darkness treatment initiation, that is, 4 h or 8 h before observation under the microscope (control cells were incubated for 4 h).

For Concanamycin A (Santa Cruz Biotechnology sc-202111, stock solution diluted in DMSO) experiments in nitrogen starvation treatment, 0,5 μM of the drug were added to solid growth media before it solidified. Protonemata were transferred to this condition 16 h previous observation under the microscope (for the 8 h deficiency time point, tissue was only incubated in Concanamycin A for 8 h).

### Chlorophyll determination

For chlorophyll content determination, chlorophyll was extracted from harvested samples with 80% (v/v) ethanol and quantified spectrophotometrically at 664 nm and 648 nm, followed by calculation using the previously established formula: c_a_ (μg/ml)= 13.36. A_664_ - 5.19.A_648_ and c_b_ (μg/ml)= 27.43 A_648_ - 8.12.A_648_.

Cell length measurements were performed on the three-first subapical cells of 7-day-old protonema grown in BCD media, stained with 10 mg. ml^-1^ of Caclcofluor, (Fluorescence brightener 28, Sigma-Aldrich). Calcofluor fluorescence was imaged with a UV filter set. For cell growth rate measurements, protonemal tissue was grown on Petri dishes (30-mm diameter) on BCD media for 7-days. Images of apical chloronemal or caulonemal cells were acquired every 10-15 min during a 2-3-h period. Protonemata cell images were taken with an Olympus BX-61 microscope equipped with a color camera (Olympus DP71). The measurements of cell length and growth rate were made using IMAGEJ software (http://rsbweb.nih.gov/ij/).

### Cell death measurements

Cell death measurement was performed according to (Castro *et al*., 2016). Briefly, moss colonies were incubated with 0.05% Evans Blue and after 2 hours tissues were washed 4 times with deionized water to remove excess and unbound dye. Dye bound to dead cells was solubilized in 50% methanol with 1% SDS for 45 min at 50°C and the absorbance was measured at 600 nm. Each biological sample consisted of 3 colonies incubated in 2 mL of the mixture methanol/SDS. Three samples were analyzed per experiment and expressed as OD/g dry weight. Dry weight was measured after drying plant colonies for 18 hours at 65°C.

### Gene expression analysis by real-time qRT–PCR

Extraction of plant total RNA was carried out using Quick-Zol Reagent (Kalium Technologies) following the manufacturer’s instructions from 7-day-old ground protonemata grown under the conditions mentioned above. 2 μg of total RNA was treated with 2 units of DNase I (Thermo Fisher Scientific) and then used as a template for cDNA synthesis with RevertAid Transcriptase (Thermo Fisher Scientific) and oligo-dT as primer following the manufacturer’s instructions. qRT-PCR was performed in thermocycler iQ5 Real-Time PCR detection system (Bio-Rad Life Science) using IQ SYBR Green SuperMix (Bio-Rad Life Science), according to the manufacturer’s instruction. The sequences of primers for qRT-PCR are described in Table S3. The genes encoding Actin and Ade-PRT were used as controls for the normalization of mRNA content between samples, according to previously published data (Le Bail *et al*., 2013). Melting curves were obtained to ensure that only a single product was amplified. The relative quantification of the target gene was obtained using Pfaffl method (Pfaffl, 2001).

### GFP-ATG8 Cleavage Assay

For the GFP-ATG8a cleavage assay, 7-day old moss protonemata proteins with the indicated treatments were harvested at the indicated time points were extracted in 100 mM Tris-HCl, 1 mM EDTA, 2% SDS, 100 mM NaCl, 1 mM PMSF and 1 μM E-64D in a relation 4:1 (buffer: fresh weight tissue). Protein concentration was measured by the Lowry method (Lowry *et al*., 1951). Equal amounts of total proteins (35 μg) were subjected to 12% SDS–PAGE and to ensure equal loading of protein samples, blots were stained with 0.5% Ponceau Red. Membranes were blocked in Tris-buffered saline (TBS; 20 mM Tris-HCl, 150 mM NaCl, pH7.4) containing 3% (weight in volume, w/v) skimmed milk powder and 0.2% Tween20 for 1h at room temperature, and then incubated with anti-GFP antibodies (Rat monoclonal, ChromoTek) diluted 1/1500 in TBS containing 0.1% Tween overnight at 4C. Anti-Mouse IgG (whole molecule)−Alkaline Phosphatase antibody (Sigma-Aldrich, A3562) diluted 1/5000 was used as the secondary antibody. The autophagic flux was expressed as the relative ratio of free GFP to GFP-ATG8a fusion quantified by ImageJ software in the same sample.

### Live-cell microscopy of GFP-ATG8-Labeled Autophagic vesicles and image analysis

Protonemata cells were observed using a Nikon Eclipse Ti confocal microscope. GFP was excited using an Ar488 nm laser and emission was collected at 512/30 nm in the EZ-C1 3.91 software. To include the whole cellular volume, z-stacks of every cell were constructed acquiring 4-5 confocal planes and compiling them with the Max. Intensity method in Fiji-ImageJ. Cell area measurements and particles quantification were also performed using Fiji-ImageJ.

### Quantification of phytohormones by LC-MS/MS

The levels of indole acetic acid (IAA) and salicylic acid (SA) in the aerial portion of plants were quantified. The extraction was carried out according to the method by (Otaiza-González *et al*., 2022), with some modifications. Briefly, 50-100 mg of tissue previously pulverized with liquid N_2_ were weighed, homogenized with 500 μL of 1-propanol/H_2_O/concentrated HCl (2:1:0.002; v/v/v), and stirred for 30 minutes at 4 °C. Then, 1 mL of dichloromethane (CH_2_Cl_2_) was applied, stirred for 30 min at 4 °C, and centrifuged at 13,000 g for 5 min. The lower organic phase (approx. 1 mL) was collected in vials, which were evaporated in a gaseous N_2_ sequence. Finally, it was re-dissolved with 0.25 mL of 50 % methanol (HPLC grade) with 0.1% CH_2_O_2_ and 50 % water with 0.1% CH2O2, and stirred slightly with vortex.

The LC-MS Waters Xevo TQs Micro (Waters, Milford, MA, USA) was equipped with a quaternary pump (Acquity UPLC H-Class, Waters), autosampler (Acquity UPLC H-Class, Waters), and a reversed-phase column (C18 Waters BEH 1.7 µm, 2.1 x 50 mm, Waters). It was used as a mobile solvent system composed of water with 0.1% CH_2_O_2_ (A) and MeOH with 0.1% CH_2_O_2_ (B), with a correction flow of 0.25 mL/min. The initial gradient of B was maintained at 40% for 0.5 min, and then linearly increased to 100% at 3 min. For identification and quantification purposes, a mass spectrometer Xevo TQ-S micro from Waters (Milford, MA, USA) coupled to the above-mentioned UPLC (LC-MS / MS) was used. The ionization source was used with electrospray (ESI), and the MassLynx Software (version 4.1) was used for data acquisition and processing. The mass spectra of the data are recorded in positive mode. The mass/charge ratio (m/z) for each metabolite were: SA: 137.0 > 65.0 and 137.0 > 93.0; IAA: 176.0 > 103.0 and 176.0 > 130.0; ABA: 263.0 > 153.0 and 263.0 > 219.0. The quantification of the activity was conducted following the calibration curves with the linear adjustment, obtaining the results in a nanogram phytohormone/milligram of fresh weight.

## RESULTS

### Differential responses of *P. patens* to carbon and nitrogen starvation

To study responses to nutrient deficiencies, known to trigger autophagy and senescence in *P. patens*, colonies of wild-type *P. patens* were grown for 14 days under optimal growth conditions (BCDAT media) and then kept under optimal growth and light conditions (LD photoperiod) either with or without nitrogen supply (nitrogen starvation) or transferred to darkness (carbon starvation) or darkness without nitrogen (carbon and nitrogen starvation) for additional 14 days. As shown in Fig. 1, *P. patens* responded differentially to the mentioned treatments. Darkness with nitrogen (DN) or without nitrogen (D-N) induced senescence and cessation of growth and development, observed by the yellowed colony phenotype (Fig. 1A), reduced values of maximum quantum efficiency PSII (Fv/Fm) (Fig. 1C), reduced growth and colony fresh weight (Fig. 1D-E), and a low number of developed gametophores (Fig. 1F) in comparison to optimal growth conditions (LN). However, after 14 days in a nitrogen-deficient medium (L-N) no senescence symptoms (Fig. 1B) or cessation of growth were observed, but rather a change in *P. patens* developmental pattern (Fig. 1B-F). Five days after transferring to L-N conditions, more caulonemata were developed (Fig. 1B upper panel), and on day 12 the gametophores exhibited longer and darker rhizoids with a reduced number of phyllids (Fig. 1B lower panel), in comparison to the LN condition. These results indicate that darkness leads to rapid senescence, whereas under nitrogen deficiency other physiological and developmental strategies, known as phenotypic plasticity, are elicited in *P. patens* allowing growth and survival for longer periods.

**Figure 1.**
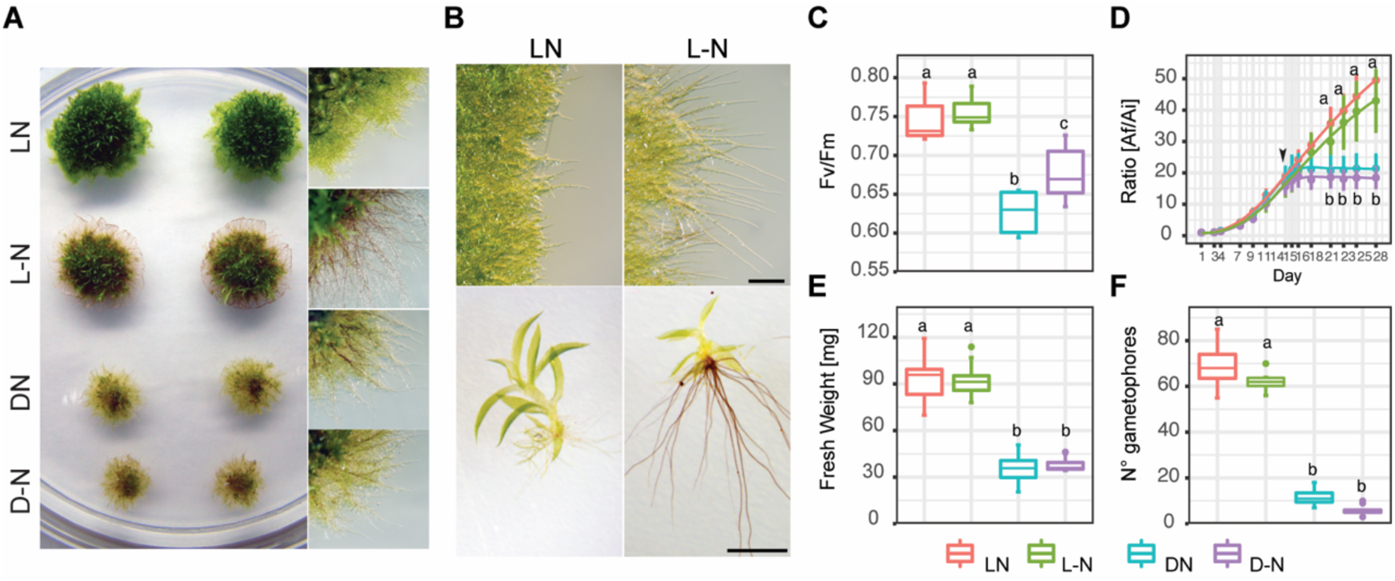
Effect of carbon and nitrogen deficiencies during a 14 day-period in colonies of *P. patens Grandsen*. **(A)** Representative images of *P. patens* colonies grown for 14 days in BCDAT (control media) and then subjected for 14 additional days to optimal growth conditions of light and nitrogen (LN), nitrogen deficiency (L-N), darkness (DN), or darkness and nitrogen deficiency (D-N). Colony view (left) and protonemata amplification view (right). **(B)** Representative images of protonemata after 5 days (upper panel) and gametophores after 12 days (lower panel) of treatment to L-N or control LN. Scale bar, 1 mm. **(C)** Box plot of PSII quantum yield (Fv/Fm), *n*=6. **(D)** Quantification of plant growth during the 28 day-old experiment was calculated as the ratio between the colony area at each time point (Af) and the colony initial area (Ai). The arrow indicates the time of transfer to deficiency media, *n*=12. **(E)** Quantification of colony fresh weight (mg), *n*=12. **(F)** Quantification of gametophore number per colony, at the end of the treatment period, *n*=6. Values represent the mean±s.d. of biological replicates. Letters indicate groups with significantly different means as determined by One-way ANOVA with Tukey’s HSD post hoc test.

### The *atg5* and *atg7* mutants exhibit both, impaired growth and development under optimal growth conditions and hypersensibility to nutrient-deficient conditions

In order to study the functional role of specific autophagic genes (*ATG*) under nutrient stress, a search for *ATG* orthologs genes was performed in the *P. patens* genome using *A. thaliana ATG* genes as a query. All core autophagy genes are present in *P. patens* (Table S1), and specific gene families such as the *PpATG8* and *PpATG18* contain several members (6 genes each), different from what is observed in *M. polymorpha* which harbors 2 *ATG8s*, 4 *ATG18s* and one gene for the other core autophagy machinery components (Fig. 2A, Table S2, (Norizuki *et al*., 2019). All *PpATG8s* belong to the Clade I *ATG8s* (Fig. S1), with the absence of members in Clade II.

**Figure 2.**
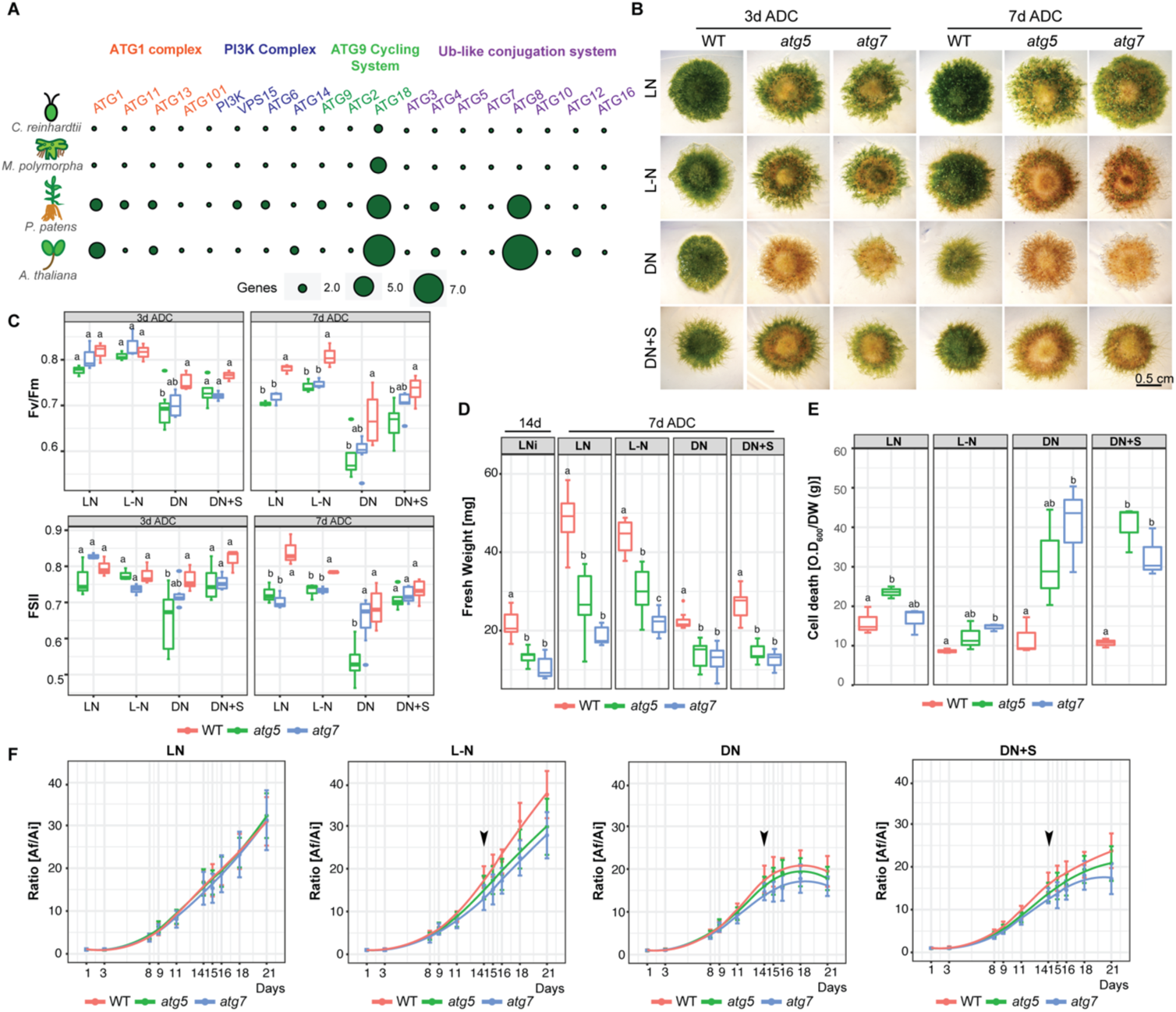
*P. patens atg5* and *atg7* mutants are hypersensitive to carbon and nitrogen starvation. (**A**) Comparison of the core autophagy genes (*ATG*) of *C. reinhardtii*, *M. polymorpha*, *P. patens,* and *A. thaliana.* **(B)** Representative images of *P. patens* colonies grown for 14 days in BCDAT (control media) and then subjected for 3 to 7 days to optimal growth conditions of light and nitrogen (LN), or nutrient deficient conditions (After Deficient Conditions, ADC) namely, nitrogen deficiency (L-N), darkness (DN), or darkness supplemented with 2% sucrose (DN+S). Scale bar, 5 mm. **(C)** Photosynthetic parameters Fv/Fm and (φPSII) of wild type, *atg5,* and *atg7* mutants at 3 days ADC or 7 days, *n*= 9. **(D)** Quantification of colony fresh weight (mg) after 14d in optimal growth conditions (LN) and at the end of the experiment, at day 21, *n*= 9. **(E)** Quantification of colony cell death with Evans Blue 7 days ADC *n*= 3. **(F)** Quantification of plant growth during the 21 day-old experiment was calculated as the ratio between the colony area at each time point (Af) and the initial colony area (Ai) *n*= 9. The arrow indicates the time of transfer to deficiency media. Values represent the mean ± s.d. of the biological replicates. Letters indicate groups with significantly different means as determined by ANOVA with Tukey’s HSD post hoc test.

It is well known that ATG5 and ATG7 are indispensable for autophagic vesicle formation, being involved in the lipidation of ATG8 with phosphatidylethanolamine, and thus knocking out these genes results in the inhibition of autophagy in several organisms (Chung *et al*., 2010). To corroborate this in *P. patens a PpATG8b::GFP-PpATG8b* reporter line that expresses *GFP-PpATG8b* (Table S1) under the control of its endogenous promoter (Sanchez-Vera *et al*., 2017) was used to generate a null *atg5* mutant and confirmed that the autophagic flux was indeed inhibited both under optimal growth conditions or carbon starvation (Fig. S2). Therefore, *atg5* and *atg7* null mutants were generated in *P. patens* by homologous recombination using targeted gene disruption in a wild-type background (Fig. S3, Table S3).

The phenotype of *atg* lines was evaluated under carbon and nitrogen starvation conditions. After 3 days of darkness, *atg5* and *atg7* lines showed accelerated senescence characterized by premature chlorosis (Fig. 2B), decreased values of the photochemical efficiency of PSII (ϕPSII), and Fv/Fm indicating damage of the photosynthetic apparatus (Fig. 2C). The phenotype observed in darkness was alleviated but not reverted when sucrose was added as an external carbon source (Fig. 2B). On the other hand, after 3 days of nitrogen deficiency (L-N), *atg5* and *atg7* phenotypes did not differ significantly from the control treatment (LN), but both *atg* lines exhibited senescence at the center of the colony in comparison to the wild-type (Fig. 2B). After 7 days in all nutrient-deficient conditions both mutant lines were affected, being the treatment of darkness the most severe followed by darkness with sucrose and then nitrogen starvation (Fig. 2B), resulting in a decline of the Fv/Fm and ϕPSII values (Fig. 2C). Remarkable to mention is that *atg* mutants maintained the nitrogen starvation response as previously observed in the wild-type, characterized by triggering caulonemata development although few and less-developed gametophores with shorter rhizoids were observed (Fig. 2B, Fig. S4).

Colony expansion measured by the radial spread of the colony was evaluated, and calculated as the ratio between the colony area at each time point (Af) and the initial area (Ai) (Fig. 2F). Under optimal growth conditions (LN) no ratio differences were observed between genotypes during the 21-day time-course experiment. However, on day 14, when lines were transferred to the nitrogen-deficient conditions (L-N), the wild-type showed a higher growth rate in comparison to the mutants. A similar trend for both mutants was observed in darkness which showed the cessation of growth already 3 days after darkness (Fig. 2B), a response that was delayed when supplemented with sucrose.

Noteworthy, even under optimal growth and LD conditions, *atg* lines exhibited a clear senescent pattern in comparison to the wild-type (Fig. 2B), which was translated into significant differences in colony fresh weight (Fig. 2D, left panel), lower photosynthesis parameters (Fig. 2B), and higher cell death (Fig. 2E). Senescence and cell death between the *atg* mutants and the wild-type were observed under all conditions tested but a different pattern was clearly observed between carbon and nitrogen deficiencies (Fig. 2D and 2E).

Taken together, although *PpATG5* and *PpATG7* are not essential genes, these results suggest that under both optimal growth or nutrient-deprived conditions, autophagy has an important contribution to *P. patens* growth and development.

### Morphological plasticity triggered in *atg5* and *atg7* to sustain protonemata growth under standard growth conditions

As previously mentioned, moss *atg* mutants maintained the nitrogen starvation response as observed in the wild-type, characterized by triggering caulonemata development although few and less-developed gametophores with shorter rhizoids were observed (Fig. 2B, Fig. S3). These results prompted us to characterize these lines under standard growth conditions on BCD media, which has nitrate as a single source of nitrogen and favors the transition from chloronemata to caulonemata, and from the juvenile to the adult gametophytic phase, allowing the observation of these two distinct differentiation processes in more detail. After 14 days of growth in BCD phenotypical differences between genotypes were observed (Fig. 3A), which exacerbated after 21 days. *atg* mutants were characterized by an early senescence phenotype with low chlorophyll content (Fig. 3B) and reduced colony weight (Fig. 3C). Similar to previous observations, colony expansion did not vary between genotypes (Fig. 3D) not only in BCD nor under BCD with external addition of sucrose (2 %), auxin (1 μM NAA), or cytokinin (0.2 μM trans-Zeatin) (Fig. 3D), indicating that protonema development in the *atg* mutants is prioritized. However, *atg* mutants exhibited a reduction in protonemata cell density (Fig. 3A and Fig. S4). This effect was also observed when plants were grown in media supplemented with sucrose or auxin, conditions known to stimulate caulonemata development (Fig. 3A and S5). Although a similar number of gametophores were developed in all genotypes (Fig. 3F), those from *atg* mutants were smaller and those at the central older part of the colony senescence prematurely (Fig. 3E). Interestingly, in the presence of trans-Zeatin, a low concentration of cytokinin known to induce bud formation, a 92% or 89% bud reduction was observed for *atg5* and *atg7,* respectively in comparison to the wild-type, suggesting that *atg* mutants are insensitive to cytokinin as a bud inducer (Fig. 3E and F).

**Figure 3.**
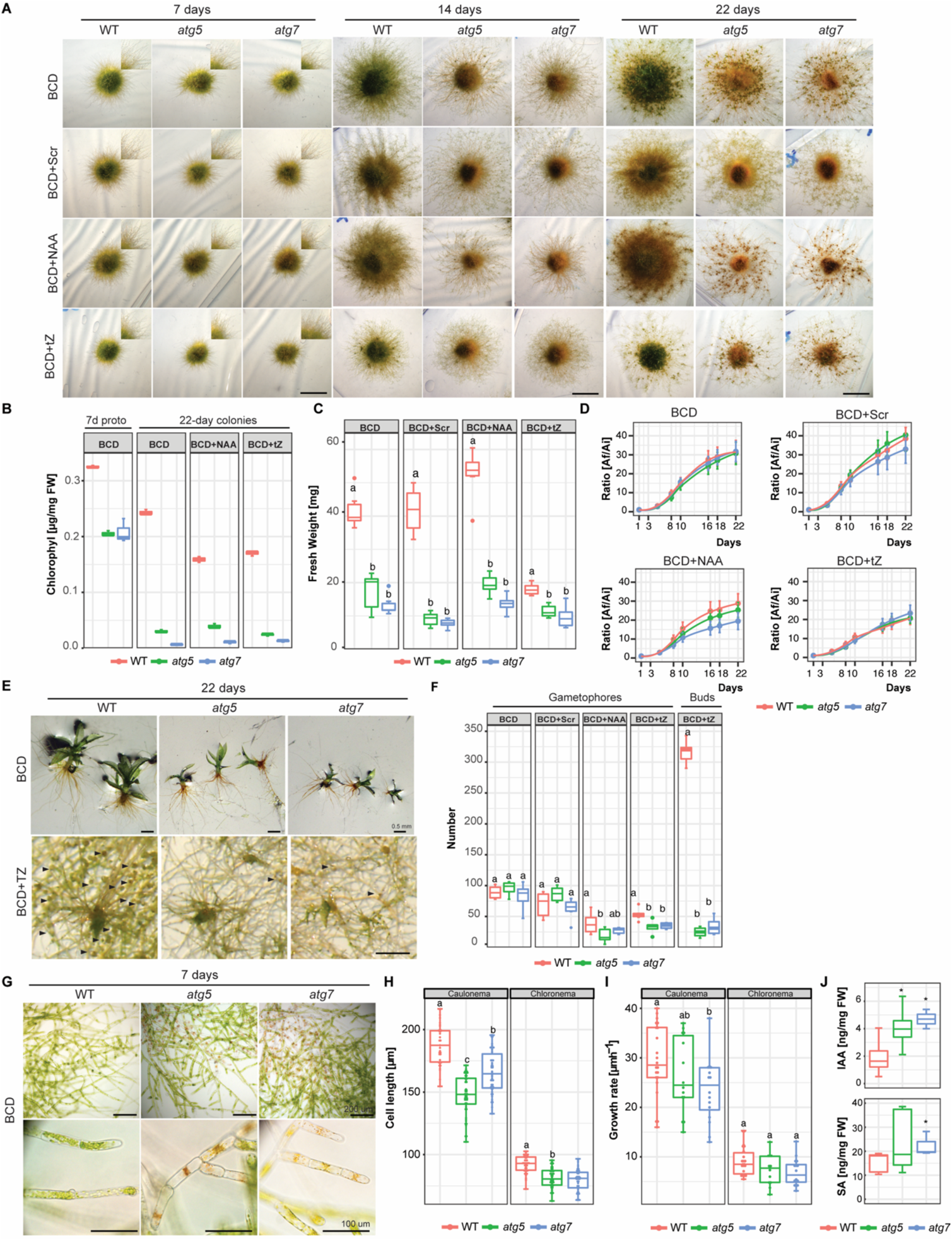
Growth and development changes of *atg* mutants in BCD media, or BCD supplemented with 2% sucrose (BCD+Scr), 1 μM 1-Naphthaleneacetic acid (BCD+NAA) or 0.2 μM trans-Zeatin (BCD+TZ). **(A)** Example images of *P. patens* colonies grown for 7, 14, and 22 days in BCD media with or without additives. Scale bar, 5 mm. **(B)** Quantification of chlorophyll (μg/mg FW) in 7-day old protonemata (starting tissue for colonies) and 22-day old colonies at the end of the experiment, *n*=3. **(C)** Quantification of colony fresh weight (mg) at the end of the experiment (day 22), *n*=6. **(D)** Quantification of plant growth during the 22 day-old experiment was calculated as the ratio between the colony area at each time point (Af) and the initial colony area (Ai), under different media, *n*=9. **(E)** Example images of *P. patens* 22*-*day old gametophores in BCD media, and buds (denoted by black arrows) induced by 0.2 μM TZ. **(F)** Quantification of gametophore number per colony or bud number after the 22-days growth period, *n*=6. **(G)** Example images of 7-day old *P. patens* protonemata. Scale bar: Upper and middle panels 200 μM, the lower panel 100 μm. **(H)** Quantification of cell length of 7-day old chloronema (*n*=44 wt, *n*= 48 *atg5*, *n*= 42 *atg7*) and caulonema cells (*n*=35 wt, *n*= 51 *atg5*, *n*= 46 *atg7*). **(I)** Quantification of the growth rate of 7-day old chloronema (*n*=22 wt, *n*= 12 *atg5*, *n*= 16 *atg7*) and caulonema cells (*n*=22 wt, *n*= 14 *atg5*, *n*= 16 *atg7*). (**J**) Quantification of auxin (Indole-3-acetic acid; IAA) and salicylic acid (SA) from wild-type, *atg5* and *atg7* mutants from 7-day old protonemata. Values represent the mean±s.d of 3 to 5 biological replicates. *, P<0.1 (by Student’s t-test, as compared with the wild type). Values represent the mean ± s.d. of the biological replicates. Letters indicate groups with significantly different means as determined by ANOVA with Tukey’s HSD post hoc test.

These protonemata differences lead us to follow thoroughly the growth of this tissue. Under standard growth conditions (BCD), senescence of the juvenile protonemata could be already distinguished in the distal older regions of 7-day-old protonemata filaments of *atg* mutants, whereas no signs of cell death were observed in the wild-type (Fig. 3G). Calcofluor staining was used to visualize cell walls and distinguish between chloronema and caulonema cells whose walls are perpendicular or oblique, respectively, to the growth axis. Both *atg5* and *atg7* lines showed a significant reduction in chloronema cell size of 12% and 15% respectively, in comparison to the wild-type, whereas a reduction of 11% and 14% in *atg5* and *atg7* respectively, was observed in caulonema cell size in comparison to the wild-type (Fig. 3H). Next, the growth rate of apical tip-growing chloronema and caulonema cells was measured and a significant reduction in caulonema apical cell growth rate but not in chloronema was observed (Fig. 3I).

The early senescence phenotype together with the protonemata growth response observed in *atg* mutants prompted us to measure the auxin (IAA), salicylic acid (SA), and abscisic acid (ABA) levels in the wild-type and *atg* mutants of 7-day old protonemata grown in BCD media by LC-MS/MS (Fig. 3J). Interestingly, both *atg* lines exhibited significantly elevated levels of IAA in comparison to the wild-type. In addition, levels for SA were higher for both *atg* mutants but only statistically significant for *atg*7, whereas ABA was not detected in the *atg* mutants nor in the wild-type (data not shown).

Overall, these results suggest that lack of autophagy triggers protonemata growth at the expense of a reduction in bud number and also gametophore development, thus suppressing or delaying the development of the adult phase of the moss life cycle. The spread of the colony is prioritized in *atg* mutants, which agrees with the investment of growth in surface area for nutrient acquisition, although cell density is diminished in addition to a reduction of protonemata cell length and caulonema cell growth rate, a cell type characterized by few chloroplasts and high growth rate in comparison to chloronemata, suggesting that autophagy contributes to sustaining the growth of this cell type.

### *PpATG8a-f* and *PpPI3KC1* genes are differentially expressed under carbon and nitrogen starvation

The early senescence pattern observed in the juvenile protonemata of *atg* mutants motivated us to further analyze the expression pattern of *P. patens* genes coding for the ubiquitin-like protein *ATG8* family (*PpATG8a-f*) and the *PI3K Complex 1* (*PpPI3K, PpVPS15a-b, PpATG6a-b, PpATG14*) involved in the early steps of autophagy induction, by real-time RT-PCR at different time points during the night of a LD photoperiod (2 h, 4 h, and 8 h), extended darkness (24 h) and during nitrogen starvation (Fig. 4A-D). The expression levels of all *PpATG8s* were rapidly induced during the first 2 h of darkness, peaked after 4 h of darkness, and by the end of the night (8 h of darkness) declined close to the levels observed at the first 2 h of darkness (Fig. 4A), being the most responsive genes *PpATG8a*, *PpATG8b,* and *PpATG8e*. During darkness supplemented with sucrose (2 % w/v), all *PpATG8s* were upregulated after 2 h, except for *PpATG8a* which was upregulated at 4 h, and all *PpATG8s* were downregulated at 4 h and 8 h of darkness. Interestingly, under extended darkness (24 h) independent of the presence of an external carbon source all *PpATG8s* were strongly upregulated, although higher expression levels were observed in the absence of sucrose. Expression of *PpPI3KCI* genes exhibited an upregulation between 2 h and 4 h of darkness and similar to *PpATG8s,* gene expression declined at 8 h. The highest expression levels for *PpATG14* and *PpPI3K* were observed after 2 h of darkness, whereas those for *PpATG6a,b,* and *PpVPS15a,b* were detected at 4 h of darkness (Fig. 4B). In the case of darkness supplemented with sucrose, all *PpPI3KC1*genes were upregulated at 2 h and 4 h, except for *PpATG6a* which was downregulated at 4 h, and all genes were downregulated at 8 h of darkness. In a similar fashion to what was observed for *PpATG8s*, if darkness was extended up to 24h with or without a carbon supply, all *PpPI3KC1* genes were upregulated.

**Figure 4.**
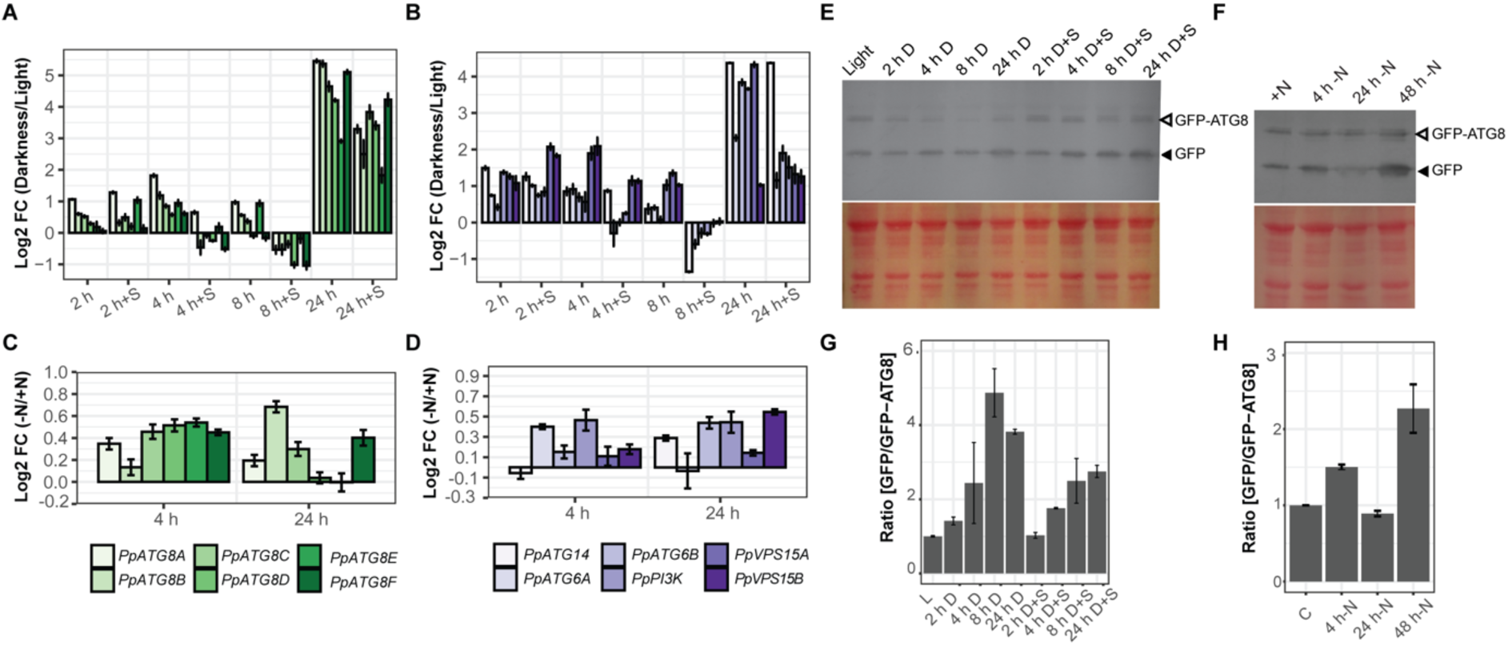
Autophagy is induced under carbon and nitrogen deficiencies. (A-D) Expression analysis of *PpATG8s* (a-f) and members of the *PI3KC* complex in *P. patens* protonemata. Log_2_-fold change values determined by qPCR upon darkness treatment or darkness supplemented with 2% Sucrose versus optimal growth conditions (light) treatment at 2h, 4h, 8h, or 24h for (**A**) *PpATG8a-f*, (**B**) *PI3KC*, or Log_2_-fold change values determined by qPCR upon nitrogen deficiency treatment versus optimal growth conditions at 2h or 24h for (**C**) *PpATG8a-f*, and (**D**) *PI3KC*. Error bars depict the standard error of the mean. (E-H) Autophagic flux analysis in *PpATG8b_pro_:GFP-PpATG8b* moss protonemata by the GFP-ATG8 processing assay. **(E)** Upper panel: Anti-GFP immunoblot was performed using protein extracts of 7-day old protonemata of *PpATG8b_pro_:GFP-PpATG8b* line under optimal growth conditions (16h light, L), or after 2h, 4h, 8h, or 24h treatment of darkness (D) or darkness supplemented with 2% Sucrose (D+S). Lower panel: Protein loading control stained with Red Ponceau. **(F)** Upper panel: Anti-GFP immunoblot performed using protein extracts of 7-day old protonemata of *PpATG8b_pro_:GFP-PpATG8b* line under continuous light and optimal nitrogen conditions (+N), or after treatment of nitrogen-deficient medium (-N) for 4h, 24h, and 48h. Lower panel: Protein loading control stained with Red Ponceau. **(G)** and **(H)** Quantification of the protein bands shown in (E) and (F), respectively. The intensity of bands corresponding to free GFP moiety (black arrowheads) normalized by the intensity of full-length GFP-ATG8 bands (grey arrowheads) is plotted (mean ± SE; n = 2).

Upon nitrogen starvation, all *PpATG8s* were slightly upregulated at 4 h and 24 h, being expression levels of all *PpATG8s* were higher at 4 h in comparison to 24 h except for *PpATG8b* which achieved the highest expression levels at 24 h (Fig. 4C). In the case of *PpPI3KCI,* all genes showed a slight upregulation at 4 h except for *PpATG14* which was upregulated at 24 h. *PpATG6a* was downregulated at 24 h, whereas *PI3K* and *VPS15a* exhibited similar induced expression levels at 4 h and 24 h (Fig. 4D).

Taken together, although both *PpATG8a-f* and *PpPI3KCI* genes were upregulated under darkness and nitrogen starvation, subtle changes in their expression signature were observed under nitrogen starvation compared to darkness. In addition, *PpATG8s* and *PpPI3KCI* expression was upregulated even with an external carbon source suggesting that the light contributes to this regulation.

A search for cis-acting regulatory elements present in the *PpATG8s* promoter regions showed that the six *PpATG8* promoters are enriched in light and stress response elements compared to the reference gene *PpADE-PRT* (Fig. S6) and five of them (*PpATG8a-e*) have a higher number of hormone response elements. Most of the elements involved in light, ABA, auxin, and abiotic stress responses are absent in the control gene, suggesting that these conditions regulate *PpATG8* gene expression in *P. patens*.

Next, a transcription factor (TF) survey of binding sites was performed on *PpATG8a-f* gene promoters using the Homer platform (Tables S4 and S5). The classes of TFs binding sites identified in the promoter region of various members from *PpATG8a-f* were those associated mainly with abiotic stress responses such as AP2/EREBP most of them belonging to the DREB, or HSF21 and HSFB3 involved in the heat shock response; with growth and development such as bHLHs and C2H2 with several associated in Arabidopsis root development, and MYB TFs involved in the regulation of circadian rhythms such as RVE7, RVE6, RVE1. Interestingly, several TFs binding sites were present in promoters from a subset of *PpATG8s*, suggesting common regulatory features (Fig. S7).

### Monitoring autophagy under carbon and nitrogen starvation through a *PpATG8b::GFP-PpATG8b* moss reporter line

Our gene expression analysis showed that *PpATG8b* (*Pp3c1_32170V3.1)* was one of the highly responsive *PpATG8* family genes to carbon and nitrogen starvation in protonemata (Fig 4A-D). Thus, to monitor autophagy induction under these conditions the *PpATG8b::GFP-PpATG8b* reporter line (Sanchez-Vera *et al*., 2017) was used to measure the autophagic flux through the GFP-ATG8 processing assay. This assay is based on the accumulation of free-GFP derived from the GFP-ATG8b fusion due to the relatively high stability of the GFP moiety once inside the vacuole, which can be detected and quantified by immunoblot analysis with anti-GFP antibodies (Chung *et al*., 2010).

Protein extracts from seven-day-old protonemata under optimal growth conditions (LD photoperiod) were collected after 16 h of light, or at different time points (2 h, 4 h, 8 h, and 24 h) during darkness or darkness supplemented with 2 % sucrose as an external carbon source. As observed in the immunoblot (Fig. 4E) and subsequent quantification (Fig. 4G), the ratio of [GFP/GFP-PpATG8b] increases during the period of darkness with the highest value at 8 h of darkness. Similarly, but to a lower extent, autophagy flux was still induced in darkness with sucrose, with increasing values as long as the dark period extended achieving similar levels at 8 and 24 h. As performed for gene expression analysis, measurements of autophagic flux under nitrogen starvation conditions were performed under continuous light, and the results indicate that autophagy flux increases at 4 h and 48 h of N starvations but decreases at 24 h (Fig. 4F and 4H).

### Autophagy is induced in apical growing cells in *P. patens*

Considering that protonema is a tissue comprising apical growing cells, the results mentioned above suggest that autophagy is involved in polar growth. To add more evidence to support this hypothesis, we first analyzed *PpATG5*, *PpATG7,* and *PpATG8a-f* expression data from a leaflet detachment experiment in which individual leaflet cells undergo reprogramming and become an apical stem cell, which consecutively generates a new protonemal filament (Busch *et al*., 2013). Reprogramming events take place between 6 h and 36 h after a leaflet is detached, matching with an increase in *PpFIE* expression, a reference gene used to indicate this developmental process. The first protonemal filaments start developing 48 h to 72 h after the incision, thus indicating the onset of apical growth events. Figure S8A shows that *PpATG8a*, *PpATG8b*, *PpATG8f*, *PpATG5,* and *PpATG7* increased their expression during the first hours after detachment, as expected for stress-induced genes and as previously reported to be a signature of autophagy during somatic reprogramming (Kanne *et al*., 2021). In addition, the expression of almost all *ATG* genes analyzed also increased in the 48 h to 72 h time window after the incision, providing more evidence that supports a link between autophagy and polar growth.

Expression data for protonemata specifically (Agudelo-Romero *et al*., 2020; Xiao *et al*., 2011) indicated that levels of most *ATG* genes are higher in caulonemata grown in optimal conditions (Fig. S8B). Considering that caulonemata elongate three times faster than chloronemata, and that these cells are very poor in chloroplasts, these observations reinforce the idea of an energetic role of autophagy in apical growth.

### Differential autophagic response in protonemata under carbon and nitrogen starvation

To get further insight into the potential connection between autophagy and polar growth, the number of autophagic vesicles in apical and subapical protonemata cells expressing *PpATG8b::GFP-PpATG8b* was quantified during the dark hours of a LD photoperiod under optimal growth conditions.

In chloronema apical cells, the number of autophagic particles significantly increases after 8 h of darkness, that is by the end of the dark hours in a LD photoperiod (Fig. 5A and 5B). However, in caulonemata, apical cells showed a significant increase in GFP-PpATG8b vesicles earlier during the night, at 4 h and 8 h (Fig. 5A and 5C). No significant differences were observed in subapical cells in any of the cases. These results suggest that autophagy could sustain the apical growth of protonemata cells during darkness under optimal growth conditions, probably acting as an energy source during the night to sustain caulonemata growth. The addition of LY294002, a PI3K inhibitor, prevented the formation of GFP-PpATG8b vesicles in light and diminished their accumulation during the night, confirming that the observed puncta are, in fact, autophagosomes (Fig. S9). The autophagic vesicles upon 24 h of extended darkness were also quantified. In this case, the number of particles remained similar to the control situation (light) in every analyzed cell (Fig. 5B and 5C).

**Figure 5.**
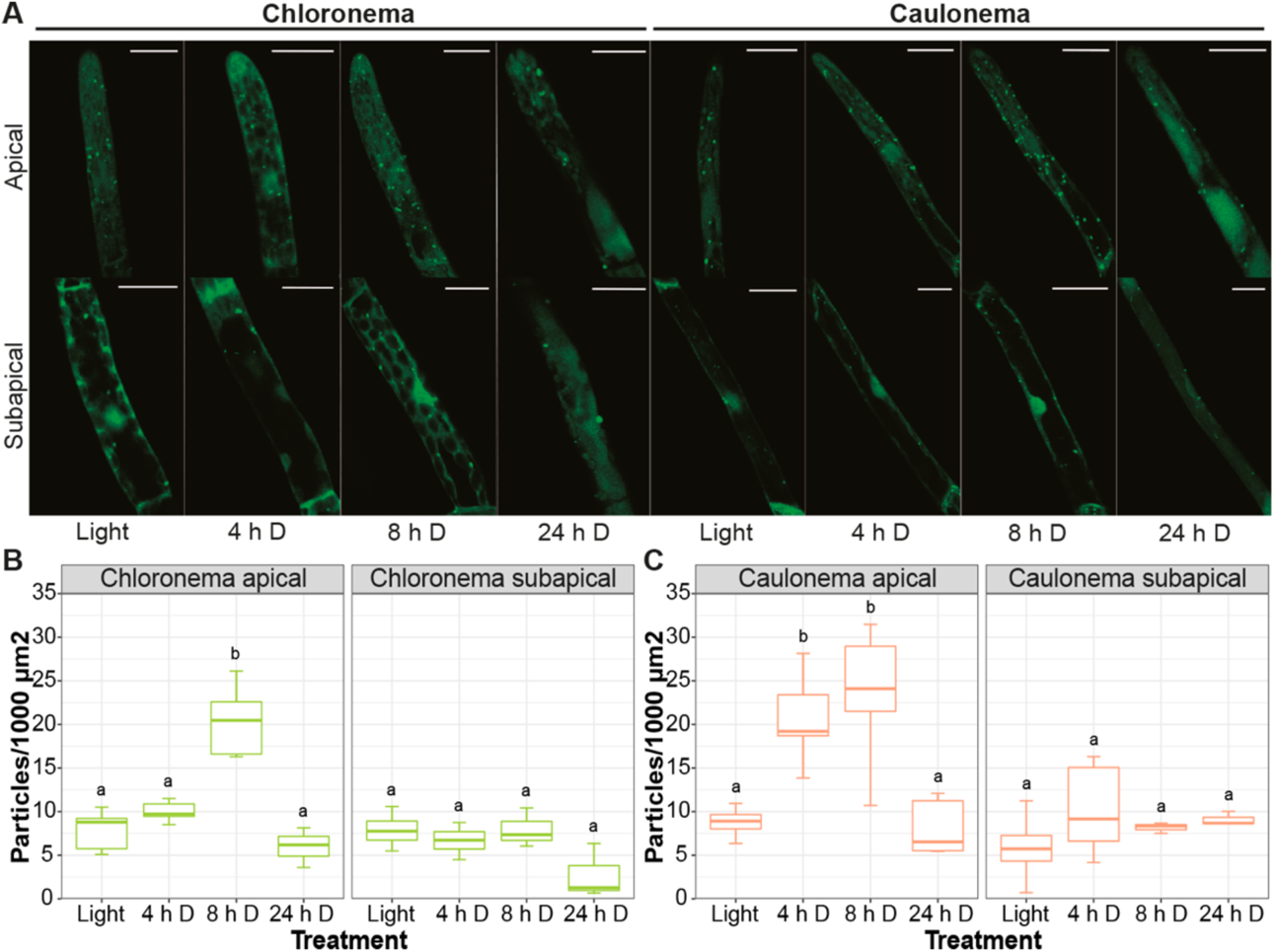
Autophagy is differentially induced in protonemata apical cells during a long day/night cycle. (**A**) Representative images of protonemata cells expressing *PpATG8b_pro_:GFP-PpATG8b* under optimal growth conditions (light, 4h and 8h darkness) and extended darkness (24h darkness). Final images were obtained from z-stacks of 4-5 confocal planes. Scale bars: 20 μm. (B-C) Quantification of autophagic particles in chloronema (**B**) and caulonema (**C**) apical and subapical cells (mean ± sd; n = 7-10). The number of particles was quantified per single cell area and then referred to 1000 μm^2^.

Next, the autophagic response of protonemata cells under nitrogen deficiency was analyzed. Concanamycin A (ConcA) is commonly used to study the accumulation of autophagic bodies under nitrogen deficiency because it reduces the acidification of the vacuole by inactivating vacuolar lytic enzymes. This is particularly relevant in experiments performed in light since darkness also triggers an inhibitory effect on H+ ATPase activity (Tamura *et al*., 2003). After exploring protonemata response to different concentrations and incubation times, cells were incubated with ConcA 0.5 μM for 16 h. Longer caulonema cells showed tip swelling and expulsion of intracellular material (Fig. 6A: caulonema, upper right panel). As DMSO-treated cells grew normally, this might be a cytotoxic effect of ConcA *per se*. Lowering ConcA concentration to 0.25 μM prevented cell damage but no significant differences were observed in treatments with or without the drug (data not shown). Due to these negative effects, quantification of autophagic vesicles under nitrogen deficit was performed without ConcA. As in Arabidopsis, in which autophagic induction by nitrogen deficiency is known to occur later than the one induced by darkness (Liu *et al*., 2020), a solid increase in GFP-PpATG8b vesicles was observed after 72 h of L-N treatment in both protonemal cell types, including apical and subapical cells (Fig. 6A-C). Figure 6 also shows that in apical cells, particles increased significantly at 8 h L-N in caulonemata and 16 h L-N in chloronemata. However, after these time points, differences with the control became slighter in both cases until 72 h of deficiency were reached. This result reinforces the idea of autophagy triggering earlier in caulonemata apical cells to sustain their growth.

**Figure 6.**
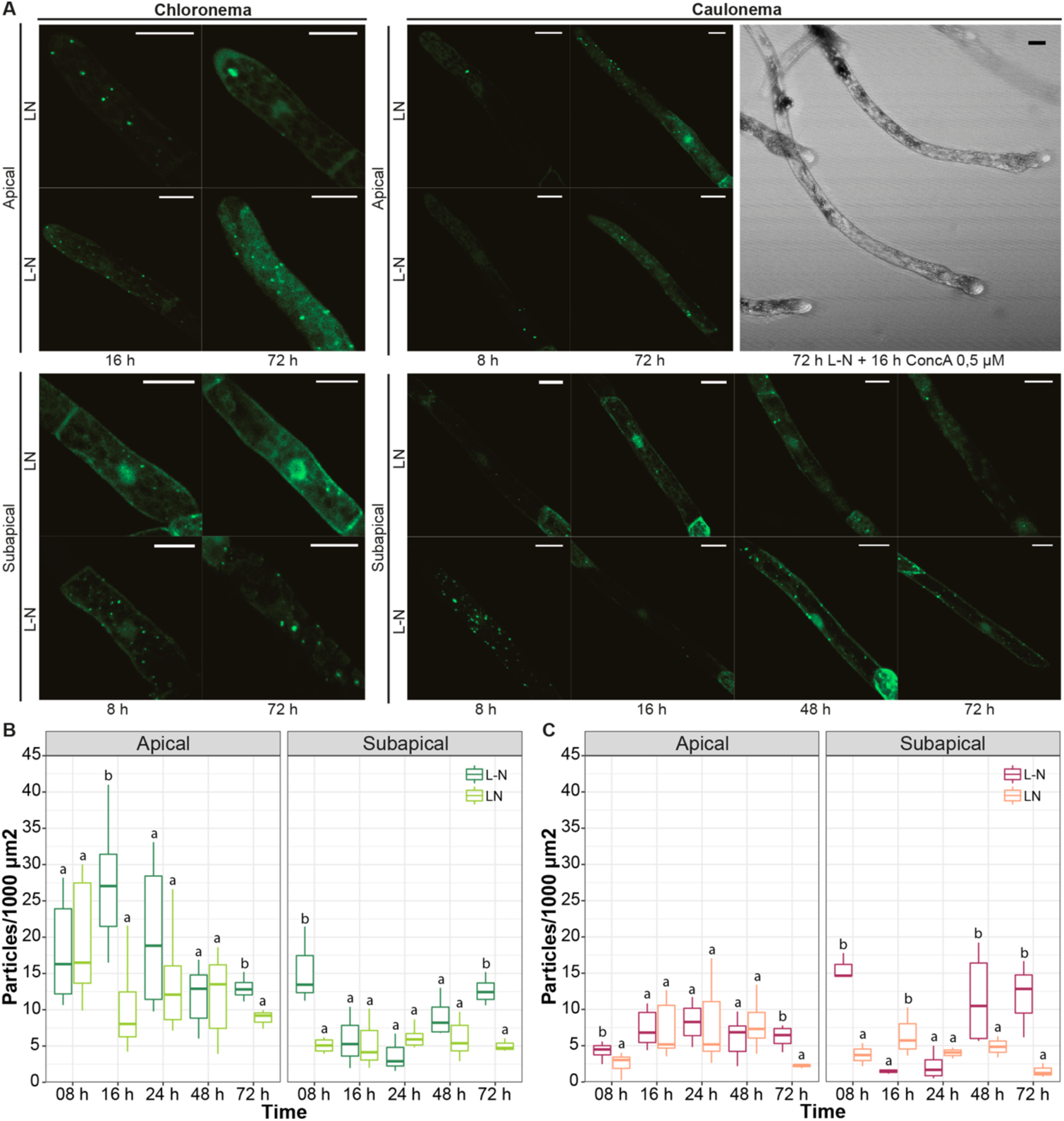
Autophagic response of protonemata cells to nitrogen deficiency. (**A**) Representative images of protonemata cells expressing *PpATG8b_pro_:GFP-PpATG8b* under optimal growth conditions (LN, BCDAT medium) and nitrogen deficiency (L-N, BCD/NaK medium; 8h, 16h, 24h, 48h, and 72h). Images shown correspond to time points that were significantly different from the controls. Cells were kept in continuous light to avoid the effect of darkness. Final images were obtained from z-stacks of 4-5 confocal planes. Scale bars: 20 μm. (B-C) Quantification of autophagic particles in chloronema (**B**) and caulonema (**C**) apical and subapical cells (mean ± sd; n = 7-10). The number of particles was quantified per single cell area and then referred to 1000 μm^2^.

A major difference between these responses and the ones induced by darkness is that subapical cells increased the number of autophagic particles under nitrogen starvation. The induction took place at 8 h and 72 h L-N in chloronemata and 8 h, 48 h, and 72 h in caulonemata. This suggests a different role for autophagy in nitrogen starvation conditions, possibly remobilizing the nutrient from subapical to apical cells to allow the latter to continue their growth.

## DISCUSSION

This study primarily aimed to determine whether autophagy plays a role in moss protonemata apical growth and hence development, in the context of the multicellular tissue of the bryophyte *P. patens*. Although it was shown that *atg5* mutants were hypersensitive to carbon and nitrogen starvation (Mukae *et al*., 2015), our work expands the knowledge on this response under both optimal and nutrient-deprived conditions through a deep phenotypic characterization of *atg5* and *atg7* mutants combined with methods to monitor autophagy. Our results provide evidence indicating that autophagy has roles during apical growth with differential responses within the cell types of the same tissue and contributes to life cycle progression and thus the development of the 2D and 3D tissues of *P. patens*.

### All core ATG proteins are present in *P. patens* with the six ubiquitin-like proteins PpATG8a-f belonging to Clade I

Our data showed that all the core *ATG* genes are present in *P. patens*. Nonetheless, specific gene families such as *PpATG8* and *PpATG18* are reduced in comparison with *A. thaliana*, but enriched compared to algae and other bryophytes such as liverworts or hornworts (Norizuki *et al*., 2019), which is probably the result of two ancestral whole-genome duplications (WGD) events in *P. patens* (Lang *et al*., 2018). Different from previous reports, our search for *ATG8s* was enriched in species from Pyramimonadophyceae, Chlorophyceae, Chlorokybophyceae, Klebsormidiophyceae, Charophyceae, Zygnemophyceae algae, and bryophytes. All the algae species contain a single-copy *ATG8* except for the single-celled *Penium margaritaceum*, a member of the Zygnematophyceae, the closest relatives of land plants, which has 2 copies. Although *P. margaritaceum* has not undergone any recent WGD, it has expanded repertoires of gene families, signaling networks, and stress responses that are associated with terrestrialization, which is in consonance with the roles of autophagy in the adaptation of plants to stress (Jiao *et al*., 2020).

The phylogenetic analysis resulted in two different ATG8 clades, confirming previous reports; Clade I including most of the Viridiplantae ATG8 members including all algae and bryophyte ATG8s, and clade II exclusively found in Ferns, Gymnosperms, and Angiosperms (Bu *et al*., 2020; Kellner *et al*., 2017; Seo *et al*., 2016). Interestingly, our results showed that several Clade I ATG8s belonging to bryophytes and streptophyte algae but not chlorophyte algae lack the extra amino acid residues at the C-terminus after the glycine residue, previously found only in some Clade II ATG8s, such as AtATG8h-i, that allows interaction with the autophagosome membrane without ATG4 processing (Seo *et al*., 2016). Thus, analyses of the different Clade I and II ATG8s will be necessary to compare their requirements for ATG4 processing and to further understand their specific functions.

### The protective role of autophagy to the canonical inducers carbon and nitrogen deficiencies is conserved in P. patens

Carbon and nitrogen starvation are canonical inducers of senescence and autophagy in plant cells. Carbon starvation-induced in response to light deprivation had a more severe effect than nitrogen deficiency in the growth and development of wild-type *P. patens*, inducing a marked rapid senescence phenotype characterized by a visible yellowing due to damage of the photosynthetic apparatus and chlorophyll degradation leading to the cessation of protonemata growth and gametophore development. Similar physiological responses to dark-induced senescence were widely described in many plants (Buchanan-Wollaston *et al*., 2005; Liebsch and Keech, 2016; Paluch-Lubawa *et al*., 2021). Moss *atg5* and *atg7* mutants showed accelerated senescence, which was mainly exacerbated under carbon starvation, where chlorophylls, photosynthesis, and weight were decreased and cell death increased, drastically in comparison to wild-type. The accelerated senescence of *A. thaliana atg* mutants has been related to the increment in ROS generation and SA levels (Masclaux-Daubresse *et al*., 2014; Yoshimoto *et al*., 2009), and an increase in SA was observed in *P. patens* although only statistically significant in *atg7*.

The symptoms caused by nitrogen deficiency in wild-type *P. patens* were more related to the induction of starvation avoidance and survival strategies, promoting the growth of caulonemata and rhizoids, and these strategies were not abolished in the *atg* mutants. Moss rhizoids though multicellular are analogous to root hairs, apical growing cells providing nutrition and anchorage to the substrate (Menand *et al*., 2007b). The stimulation of rhizoid growth of wild-type *P. patens* under nitrogen deficiency resembles Arabidopsis root hair growth triggered under low-nitrate conditions (Vatter *et al*., 2015). Although moss *atg* mutants maintained the development of caulonemata and rhizoids, final cell sizes were shorter, and growth was sustained at the expense of the early senescence of the central part of the colony, by reducing gametophore number and size (see discussion below).

### Autophagy contributes to *P. patens* growth program and its absence triggers changes in development prioritizing protonemata growth at the expense of the adult gametophytic phase

Arabidopsis *atg* mutants, with exception of those genes involved in the PI3K complex (PI3K, VPS15, ATG6), when grown under optimal growth conditions and LD photoperiod germinated and developed as wild-type plants (Izumi *et al*., 2013). However, under these conditions, *P. patens atg5* and *atg7* mutants exhibited several defects in growth and development in gametophores and protonemata, such as stunted gametophores, and premature senescence, hyposensitivity to cytokinin, and a dampened response to sucrose and auxin, which are known to stimulate caulonemata growth. Moreover, *atg5* and *atg7* did not show defects to differentiate chloronemal and caulonemal tip growing cells, albeit differences such as reduced chloronema and caulonema cell length and reduced apical caulonemata growth rate were observed, suggesting that autophagy has a role sustaining their apical growth. The role of autophagy in polar growth in plants has almost been underexplored, with a study showing that PTEN, a phosphatase of PI3P regulates autophagy in pollen tubes (Zhang *et al*., 2011), but it is known to have a role in other organisms such as fungi. Caulonema tip growth is reminiscent of fungal hyphae (Bartoszewska and Kiel, 2011) since the *atg5* and *atg7* caulonemata growth could be compared with the response of the *atg1* line of *Aspergillus fumigatus* which, also resulted in inhibition of colony growth under starvation (Richie *et al*., 2007). These facts revealed the physiological importance of autophagy for polar tip growth of filamentous tissue in charge of nutrient acquisition and its adaptive role to cope with starvation (Richie *et al*., 2007). Interestingly, new evidence has provided insights supporting the notion that autophagy may have effects on vacuolar enlargement, which in turn could impact on cell elongation (Robert *et al*., 2021a; Robert *et al*., 2021b).

Nevertheless, under all conditions tested, it was revealed a clear strategy of *atg* mutants at the whole plant level, where protonemata growth is prioritized at the expense of gametophore development and thus life cycle progression. The higher levels of auxin measured in protonemata in *atg* mutants suggest that the signal to trigger the differentiation of more caulonemata cells is present, which was not translated into an increase of this cell type, probably preventing unaffordable growth according to the moss energetic resources. Other mutants impaired in energy signaling such as the double knock-out of trehalose-6-phosphate synthase (*tps1-tps2*), and the hexokinase-1 (*hxk1*), are negatively affected in caulonemata differentiation (Olsson *et al*., 2003; Phan *et al*., 2020). Moreover, the phenotypes of the energy sensor Snf1-related protein kinase 1 (*snf1a-snf1b*), known to activate autophagy and inhibit the TOR-kinase complex in Arabidopsis (Soto-Burgos and Bassham, 2017), shared several phenotypes in common with the moss *atg5* and *atg7* lines such as premature senescence, few leafy shoots with shorter stems and smaller leaves, hyposensitivity to cytokinins, and shorter chloronemata cells (Thelander *et al*., 2004). However, the *snf1a-snf1b* exhibited an excess of caulonemata due to an increased sensitivity to auxin, few aberrant chloronemal filaments, and failed to grow in a LD photoperiod (Thelander *et al*., 2004); which are severe phenotypes in comparison to the *atg* mutants possibly due to the fact that SNRK1 is upstream autophagy, and integrated signals from multiple pathways.

### Autophagy is differentially induced in moss protonemata by carbon or nitrogen starvation

Gene expression analysis under carbon starvation during the night of a LD photoperiod, and during extended darkness (24 h), with and without sucrose allowed to study the effect of light on autophagy. A general trend observed for several *PpATG8s* and *PpPI3KC1* genes was their up-regulation with the shift from 16h light to darkness during the first 4 h of darkness. At 8 h of darkness, albeit their expression levels were still up-regulated they diminished closer to the 2 h darkness time point. Similar behavior was reported for *ATG6*, *PI3K*, *ATG7*, and *ATG9* in Arabidopsis during the night, associated with leaf starch degradation by autophagy (Wang *et al*., 2013). Interestingly, our *PpATG8s* promoter analysis showed several TFBs involved in the regulation of circadian rhythms. After extended darkness (24 h) the levels of all genes were up-regulated reaching the highest levels of expression in the medium with or without sucrose, strongly suggesting the role of light in autophagy induction, consistent with previous observations in Arabidopsis showing that light signaling could regulate autophagy through modulating the transcription level of *ATGs* (Yang *et al*., 2020).

Another approach used to study the autophagic response in protonemata was the quantification of autophagic particles in the *PpATG8b::GFP-PpATG8*b reporter line. The observation of cells during the night (4 h and 8 h darkness) revealed a differential response depending on the cell type and the position of the cell along the protonemal filament. Apical cells, which are actively growing, increased the number of GFP-PpATG8b particles during the night. Interestingly, earlier increases were observed in caulonemata compared to chloronemata. On the other hand, the basal number of autophagic vesicles in subapical cells, in which growth has ceased, did not change during the dark hours of the photoperiod.

In vascular plants, autophagy contributes to the increase in energy supply through the degradation of stromal proteins into free amino acids, RCB pathway (Ishida *et al*., 2008) or starch granules breakdown, SSGL body pathway (Wang *et al*., 2013). Considering the higher growth rate and the lower number of chloroplasts in caulonema apical cells compared to chloronema, our observations support the idea of autophagy triggering earlier during the night in caulonemata as an energy-providing mechanism to sustain the growth processes in the dark. These results are consistent with the above-mentioned phenotype reflecting impaired energetic resources in *atg* mutants.

Our qPCR data and autophagic flux analysis indicate that autophagy is activated in extended darkness (24 h) but the number of GFP-PpATG8b particles appears to stay the same as in the control situation. This may be due to an enhanced autophagic flux rapidly delivering vesicles to the vacuole for degradation, or free GFP accumulation in vacuoles under dark conditions which increases vacuolar fluorescence making it difficult to distinguish autophagic bodies in the lumen.

In tissue grown in light without nitrogen (L-N), the time difference in autophagic particles accumulation between chloronema and caulonema apical cells was still observed, but a different response was detected in subapical cells compared to darkness. In this case, subapical cells of both protonemata cell types increased the number of autophagic particles at different time points. These observations are compatible with the previously described role of autophagy in nitrogen remobilization during senescence and nitrogen starvation in vascular plants (Guiboileau *et al*., 2012; Li *et al*., 2015). Our observations showed that in the *atg* mutants older cells of a filament died prematurely, suggesting that another mechanism than autophagy could be involved in recycling components of older cells to sustain protonemata apical growth. During energy starvation in Arabidopsis, an autophagic-independent route of chloroplast degradation associated with the Chloroplast vesiculation (CV) pathway is highly induced in the absence of autophagy contributing to the early senescence phenotype of *atg* mutants (Wang and Blumwald, 2014). However, we did not find homologs of the CV gene (AT2G25625) neither in algae nor in bryophytes genomes.

Taken together, our work expands the knowledge of autophagy in bryophytes, has identified several components of the *P. patens* autophagic system, and provides valuable results indicating that autophagy has roles during apical growth highlighting a degree of cell-type specificity in the autophagy response within the same tissue, which in turn contributes to the development of the 2D and 3D tissues of *P. patens*.

## SUPPLEMENTARY DATA

**Table S1**. List of *ATG* genes from *Physcomitrium patens* and *Arabidopsis thaliana*.

**Table S2**. List of ATG8 proteins across several Viridiplantae species.

**Table S3**. List of primers used in this study.

**Table S4**. Analysis of *PpATG8s* transcription factor binding sites (TFBSs) from the Homer plant database.

**Table S5**. Co-expression network table of *PpATG8s* promoters generated using “NetworkAnalyzer” tool from Cytoscape, keeping nodes with at least two degrees.

**Figure S1.** Phylogenetic tree of ATG8s.

**Figure S2**. The autophagic flux is blocked in *P. patens atg5-(PpATG8b::GFP-PpATG8b)* knock-out mutant.

**Figure S3**. Constructs used for homologous recombination and genotyping *atg5* and *atg7* knock-out mutants.

**Figure S4**. Morphological changes of *atg* mutants under nitrogen-deficient conditions.

**Figure S5.** Morphological changes of *atg* mutants under BCD media growth conditions, or BCD supplemented with sucrose, auxin or cytokinin at 14 or 22 days of growth.

**Figure S6**. Analysis of Cis Acting Regulatory Elements (CARE) identified in *PpATG8a-f* promoters.

**Figure S7**. Gene–motif interaction network of *PpATG8s* genes.

**Figure S8.** *In silico* expression analysis of several *PpATG* genes regarding apical growth

**Figure S9.** Quantification of autophagic particles in *PpATG8b::GFP-PpATG8b* chloronema cells treated with LY294002.

## Supporting information

List of ATG genes from Physcomitrium patens and Arabidopsis thaliana.

List of ATG8 proteins across several Viridiplantae species.

List of primers used in this study.

Analysis of PpATG8s transcription factor binding sites (TFBSs) from the Homer plant database.

## Acknowledgments

We are extremely grateful to Dr. Victoria Sanchez-Vera and Dr. Mattias Thelander (Swedish University of Agricultural Sciences) for the gift of the *atg5-ko* and *atg7-ko* plasmids for gene disruption in *P. patens*, and for the *PpATG8b-GFP* reporter line. We also acknowledge Dr. Alejandra Trenchi (IMBIV, CONICET-UNC) and Dr. Leandro Ortega (INTA) for technical support with laser scanning confocal microscopy. We acknowledge CONICET for the Ph.D. scholarship to G.P and Secyt-UNC for the EVC-CIN scholarship to J.F and F.L. We acknowledge Dr. Damián Cambiagno for helpful discussions.

## Author contributions

R.L and L.S conceived the project; R.L, L.S, and G.P designed the experiments; S.O-G, P.V, and M.T performed the hormonal quantification and P.A-R performed the *PpATG8s* promoter network analysis. All remaining experimental work was performed by L.S, G.P, J.F, M.P-R, and F.L. L.S, G.P, J.F, F.L, G.R, C.G and R.L analyzed the data. L.S, G.P and R.L wrote the manuscript draft and incorporated feedback from all authors.

## SUPPLEMENTAL MATERIAL

**Figure S1.**
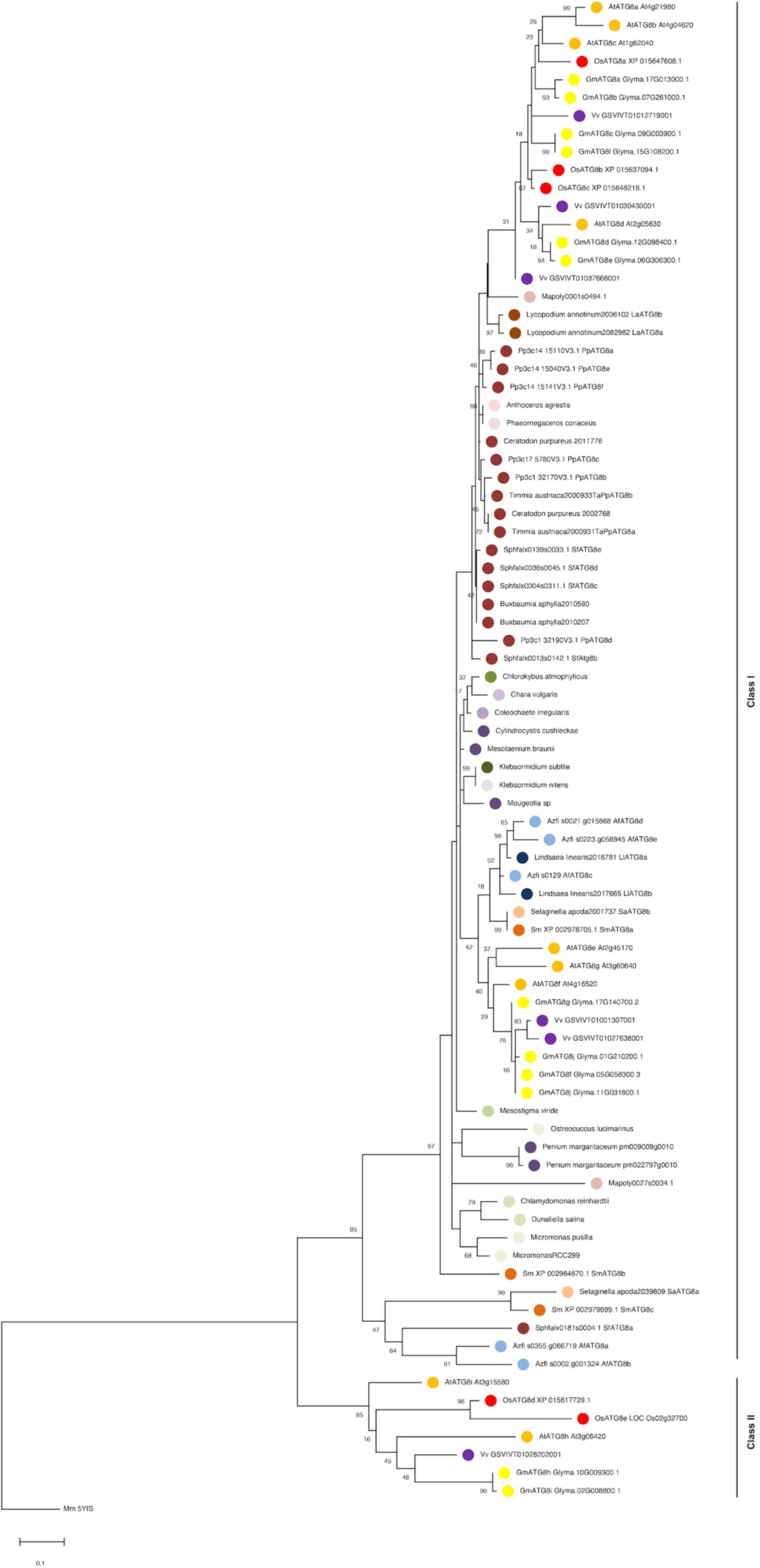
Phylogenetic tree of ATG8s. The evolutionary history was inferred by using the Maximum Likelihood method and JTT matrix-based model. The percentage of trees in which the associated taxa clustered together is shown next to the branches. The tree is drawn to scale, with branch lengths measured in the number of substitutions per site. This analysis involved 84 amino acid sequences. All positions with less than 95% site coverage were eliminated, i.e., fewer than 5% alignment gaps, missing data, and ambiguous bases were allowed at any position (partial deletion option). Evolutionary analyses were conducted in MEGA X [2].

**Figure S2.**
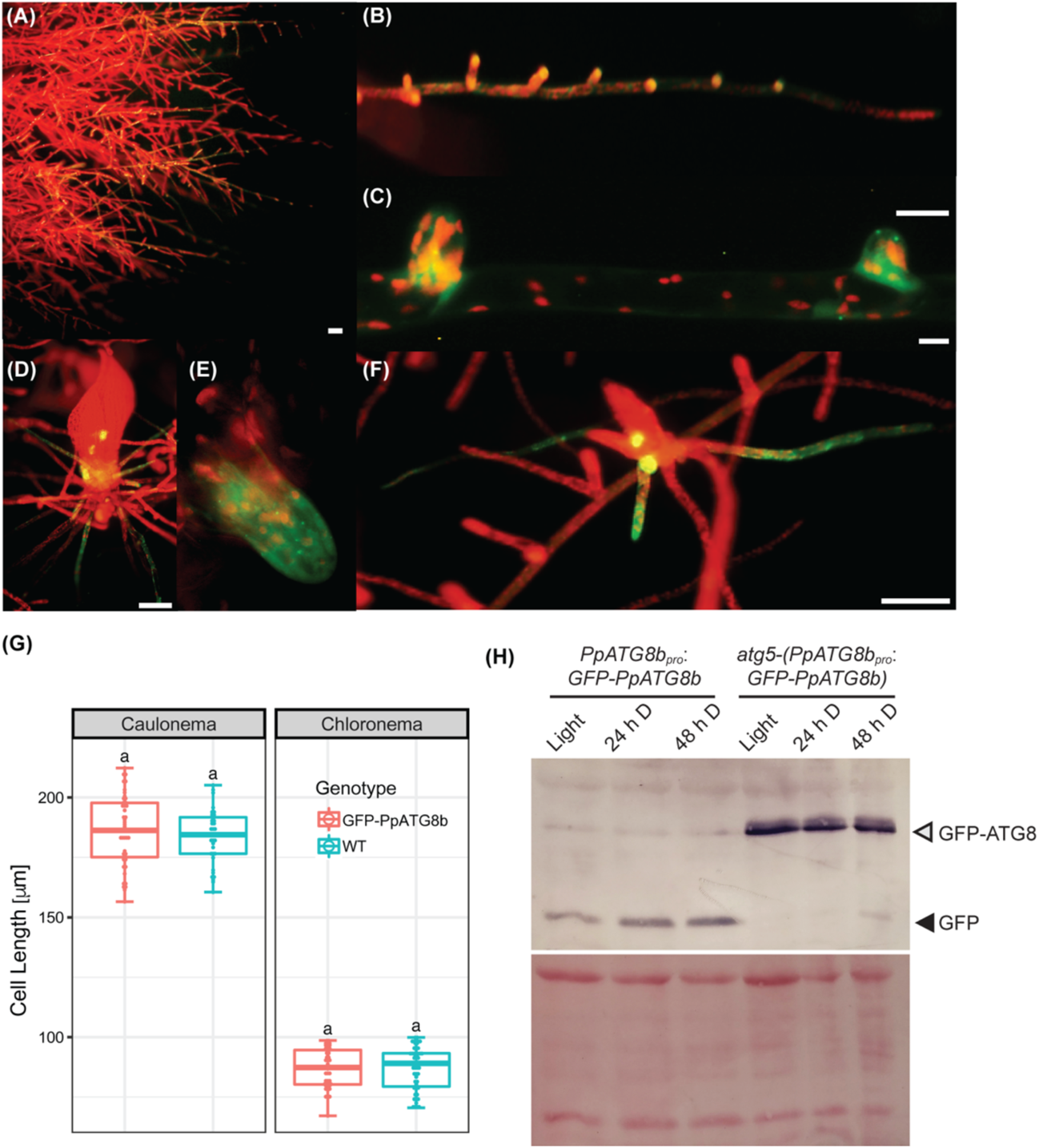
The autophagic flux is blocked in *P. patens atg5*-(*PpATG8b_pro_:GFP-PpATG8b)* loss-of-function mutants. *PpATG8b_pro_:GFP-PpATG8b* autophagy marker line: (A) Protonemata, (B) Magnification of a protonemal filament, (C) Branching, (D) Adult gametophyte, (E) Young rhizoid cell, (F) Young gametophyte. (G) Box plot of 7 d old protonemata cell size: Chloronemata and Caulonemata. Bar 500 μm. (H) Upper panel: Anti-GFP immunoblot performed using protein extracts of 7-day old protonemata of *PpATG8b_pro_:GFP-PpATG8b* or *atg5-PpATG8b_pro_:GFP-PpATG8b* line under optimal growth conditions (16 h light, L), or after 24 h or 48 h treatment of darkness. Lower panel: Protein loading control stained with Red Ponceau.

**Figure S3.**
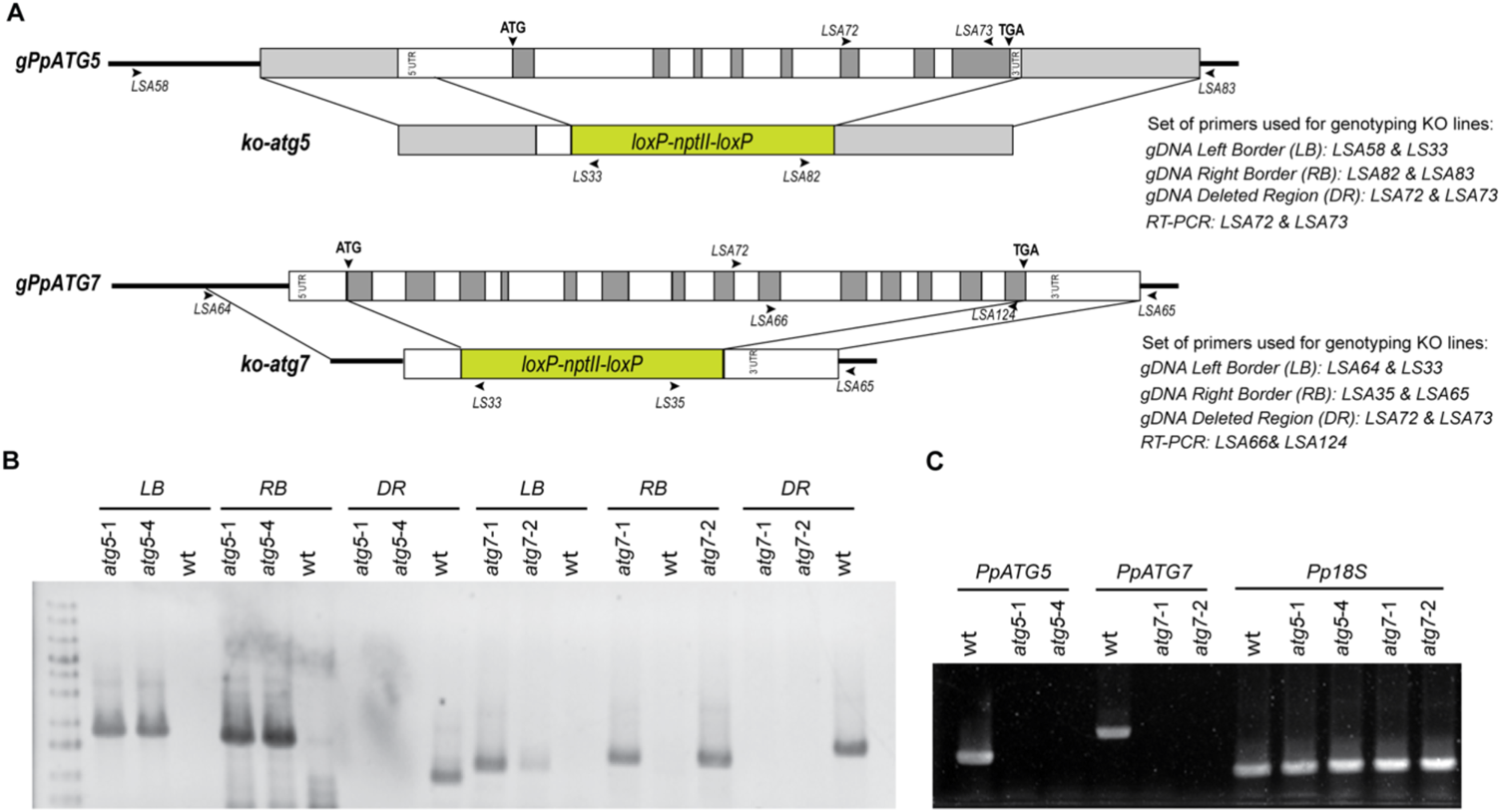
Constructs used for homologous recombination and genotyping *atg* KO plants. (**A**) Schematic representation of the genomic locus for *PpATG5* and *PpATG7*, and the derived constructs used in this study for the disruption of the gene by homologous recombination. White and grey boxes correspond to exons and introns, respectively. The locations of the primers used for genotyping are shown by arrowheads. Primers sequences are listed in Supplemental Table 2. (**B**) PCR genotyping analysis of wild-type, *atg5* and *atg7* knockout lines. The location of the primers used for genotyping is shown in A. Gene targeting events were detected by simultaneous PCR amplifications utilizing gene-specific primers external to the targeting construct in combination with outward-pointing primers specific to the selectable marker cassette. PCR genotyping with primers towards the deleted region, amplified only when the *atg5 or atg7* locus is unaltered, but not in the knockout lines. Left border (LB); right border (RB), deleted region (DR). (**C**) RT-PCR of wild-type, *atg5,* and *atg7* knockout lines showing the absence of *atg* transcripts in the knockout lines.

**Figure S4.**
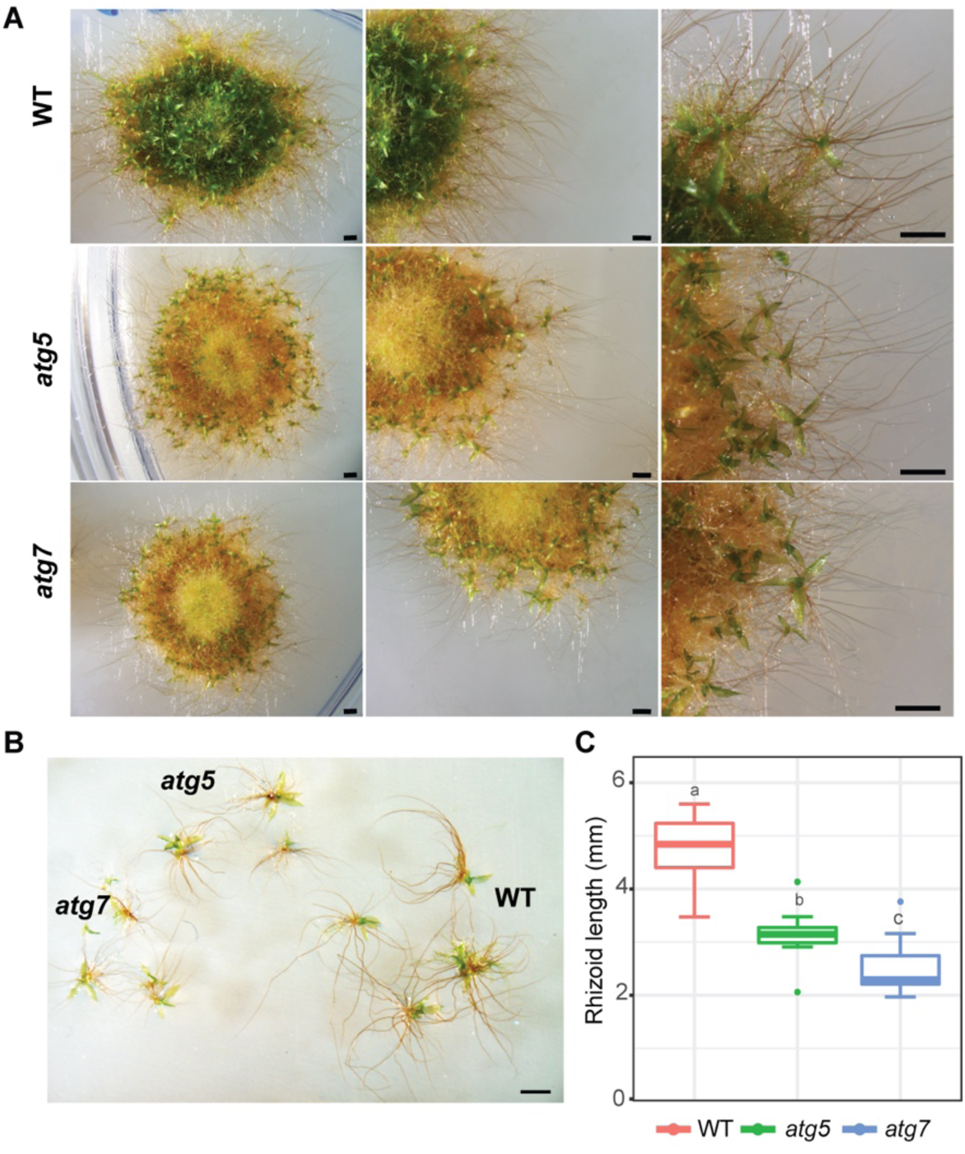
Morphological changes of *atg* mutants under nitrogen-deficient conditions. **(A)** Representative images of *P. patens* colonies grown for 14 days in BCDAT (control media) and then subjected for 7 additional days to nitrogen deficiency, Colony view (left), colony amplification view showing growing gametophores (middle), and gametophore amplification view (right). **(B)** Representative gametophores from wild-type, *atg5,* and *atg7* knockout lines. **(C)** Quantification of gametophore rhizoid length (mm) at the end of the experiment (day 22), n=8-10.

**Figure S5.**
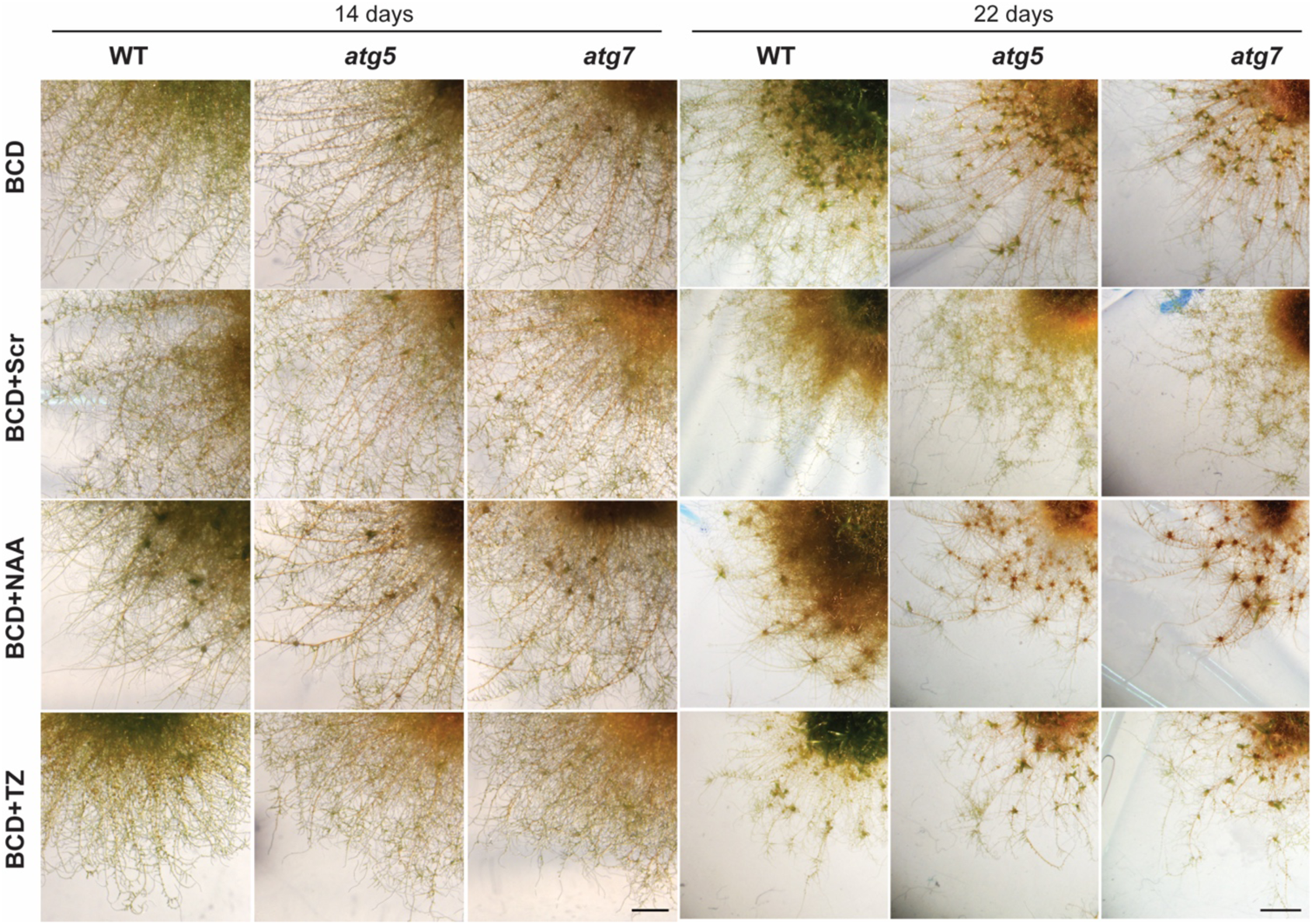
Morphological changes of *atg* mutants under BCD media growth conditions, or BCD supplemented with 2% sucrose (BCD+Scr), 1 μM 1-Naphthaleneacetic acid (BCD+NAA) or 0.2 μM trans-Zeatin (BCD+TZ) at 14 or 22 day of growth. Images show a piece of *P. patens* colonies and protonemata protruding from it. Scale bar 0.5 cm.

**Figure S6.**
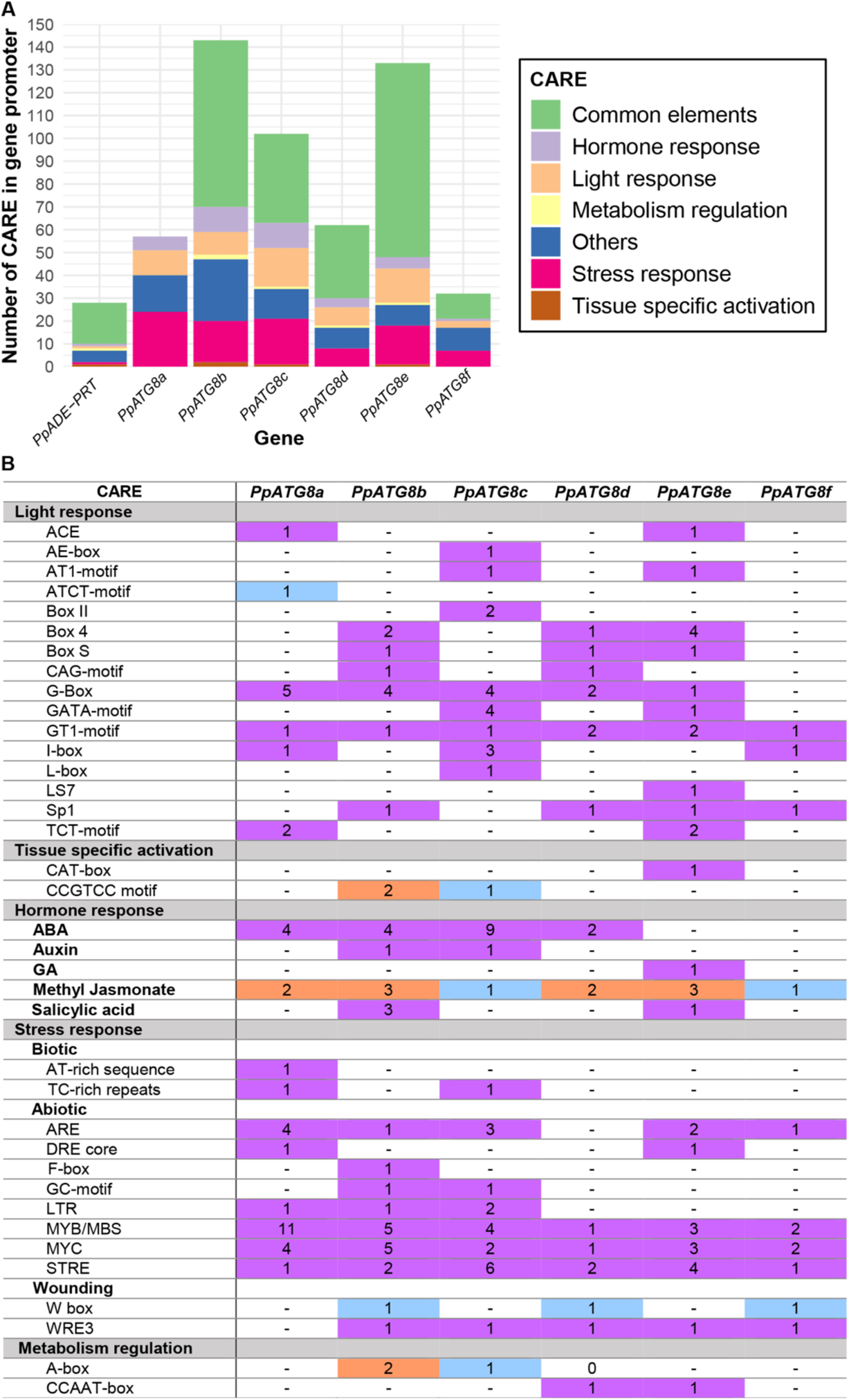
Analysis of Cis Acting Regulatory Elements (CARE) identified in *PpATG8a-f* promoters. (**A**) Comparison of CARE composition in each promoter region based on their biological functions. *PpADE-PRT* was used as a reference gene. “Common elements” category includes TATA-box and other promoter-common motifs involved in transcription regulation (activation/repression). Motifs with unknown functions are listed as “Others”. (**B**) List of the elements included in each CARE category (“Common elements” and “Others” are not taken into account). Numbers indicate how many times the element is present in the sequence. CARE not found in the reference gene are shown in light purple; CARE present in a higher number compared to the reference gene are shown in salmon; CARE present in a lower or same number compared to the reference gene are shown in light blue.

**Figure S7.**
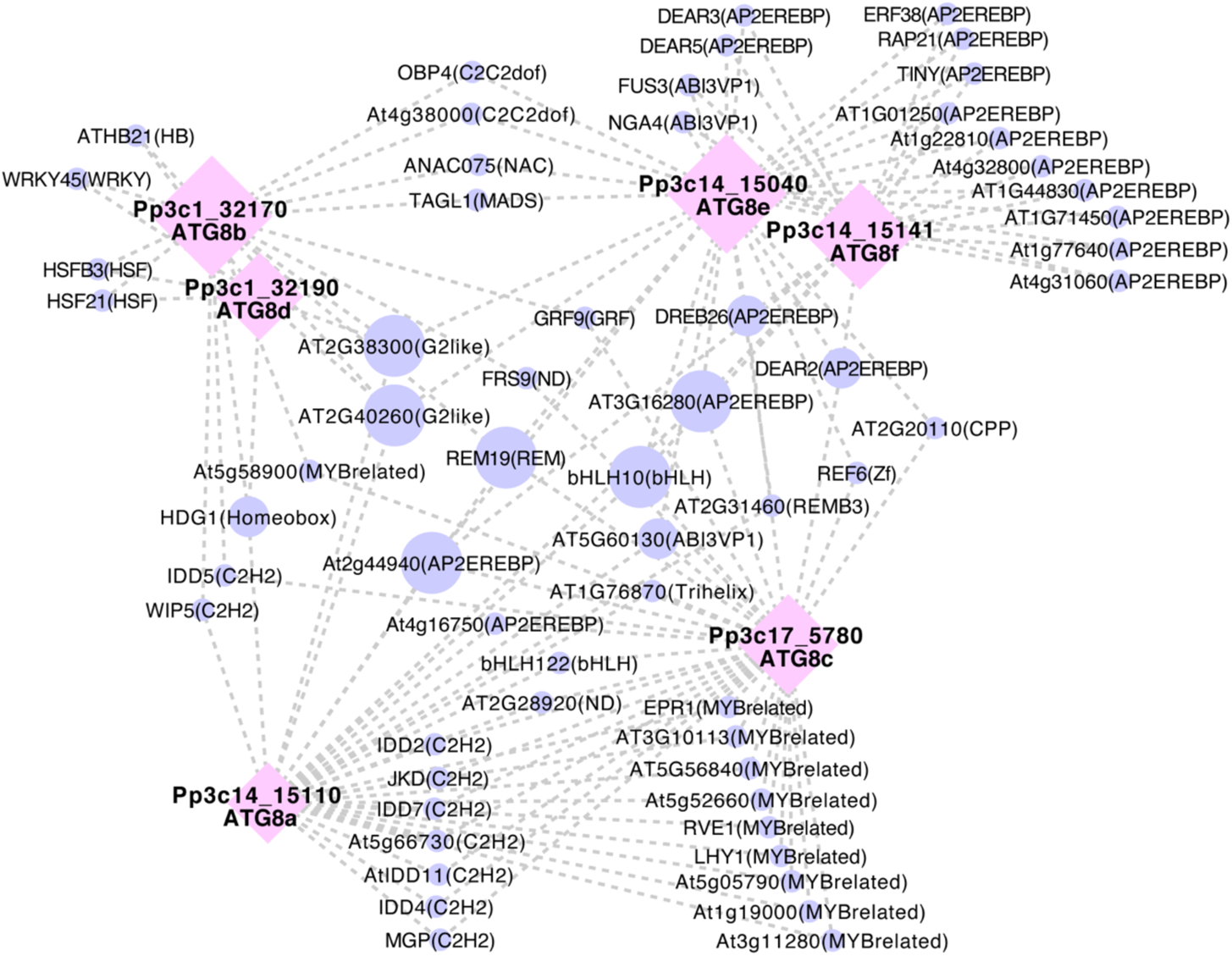
Gene–motif interaction network of *PpATG8s* genes. Motifs were filtered by a motif Score > 9 and with at least two degrees (Tables S4 and S5). Nodes are represented as *PpATG8s* transcripts (purple circle) and motifs (pink diamond), also their size displays the number of connections. Edges show their interaction (gray dash line).

**Figure S8.**
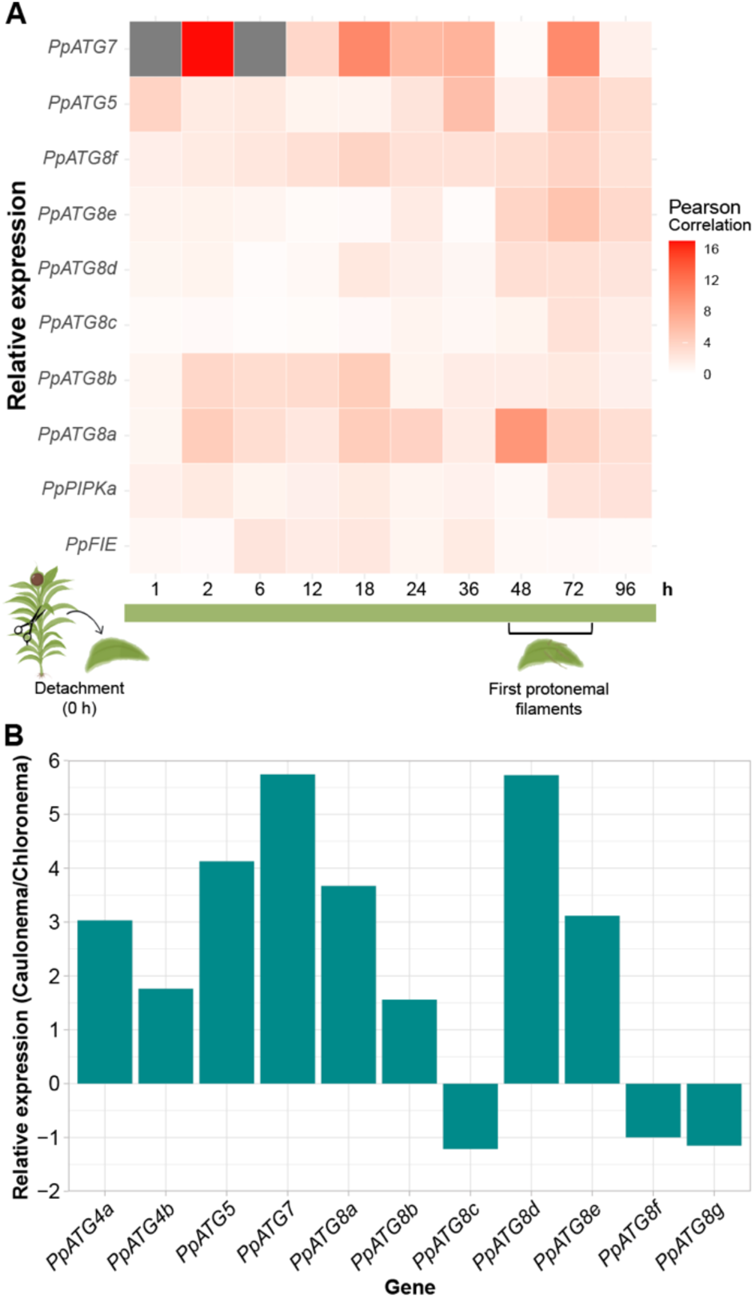
*In silico* expression analysis of several *PpATG* genes regarding apical growth. (A) Heat map of relative expression during reprogramming and apical growth in *P. patens*. *PpFIE* was used as a reference gene for reprogramming events and *PpPIPKa* as reference for apical growth processes. Values were obtained from PEATmoss database, Combimatrix Leaflet Development gmv1.2 dataset, (Fernandez-Pozo, Haas et al. 2020). (B) Relative expression of *PpATG8s* genes indicated as a caulonema to chloronema expression ratio. Data obtained from Xiao L. et al, 2011.

**Figure S9.**
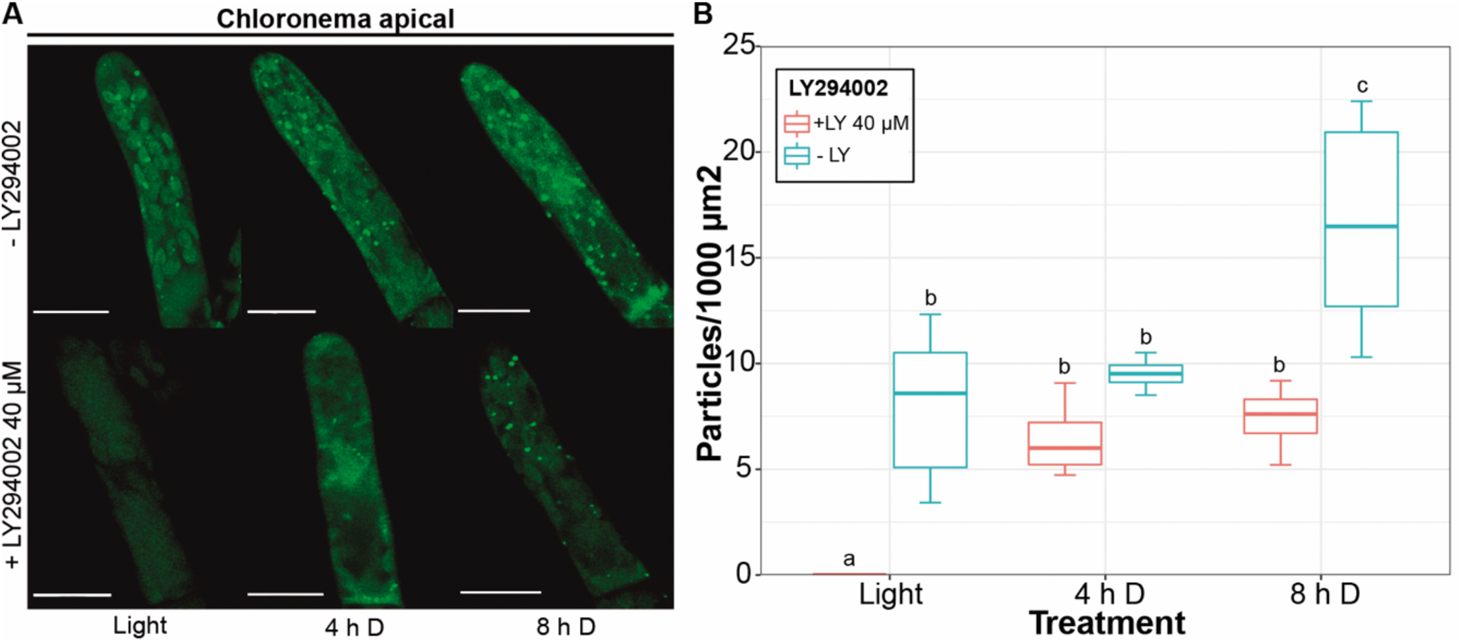
Quantification of autophagic particles in *PpATG8b_:_:GFP-PpATG8b* chloronema cells treated with LY294002 40 µM. (A) Representative images of chloronema apical cells expressing *PpATG8b_:_:GFP-PpATG8b* under optimal growth conditions (light, 4h, and 8h darkness). Final images were obtained from z-stacks of 4-5 confocal planes. Scale bars: 20 μm. (B) Quantification of autophagic particles (mean ± sd; n = 7-10). The number of particles was quantified per single cell area and then referred to 1000 μm^2^.

